# A world of viruses nested within parasites: Unraveling viral diversity within parasitic flatworms (Platyhelminthes)

**DOI:** 10.1101/2021.12.15.472606

**Authors:** Nolwenn M Dheilly, Pierrick Lucas, Yannick Blanchard, Karyna Rosario

## Abstract

Because parasites have an inextricable relationship with their host, they have the potential to serve as viral reservoirs or facilitate virus host-shifts. Yet, little is known about viruses infecting parasitic hosts except for blood-feeding arthropods that are well-known vectors of zoonotic viruses. Herein we uncover viruses of flatworms (Phylum Platyhelminthes, group Neodermata) that specialize in parasitizing vertebrates and their ancestral free-living relatives. We discovered 115 novel viral sequences, including 1 in Macrostomorpha, 5 in Polycladida, 44 in Tricladida, 1 in Monogenea, 15 in Cestoda and 49 in Trematoda, through data mining. The majority of newly identified viruses constitute novel families or genera. Phylogenetic analyses show that the virome of flatworms changed dramatically during the transition of Neodermatans to a parasitic lifestyle. Most Neodermatan viruses seem to co-diversify with their host, with the exception of rhabdoviruses which may switch host more often, based on phylogenetic relationships. Neodermatan rhabodviruses also have an ancestral position to vertebrate-associated viruses, including Lyssaviruses, suggesting that vertebrate rhabdoviruses emerged from a flatworm rhabdovirus in a parasitized host. This study reveals an extensive diversity of viruses in Platyhelminthes and highlights the need to evaluate the role of viral infection in flatworm-associated diseases.

## Introduction

Over the past few years, high-throughput sequencing technologies have expanded the virosphere well beyond pathogenic viruses and/or viruses that can be cultured. Therefore, the known taxonomy and understanding of virus evolution have dramatically increased^1^. One such breakthrough is the finding that the diversity of plant- and vertebrate-infecting RNA viruses is nested within the diversity of viruses that infect arthropods^2^ suggesting that these invertebrates have played a role as viral reservoirs and facilitated host shifts^2,3^. Despite these advances, our current knowledge of the global RNA virome remains strongly biased, with most sequencing efforts focusing on plants, chordates and arthropods, and to a lesser extent nematodes and mollusks. Sampling viruses from a larger diversity of eukaryotic hosts should lead to new and improved evolutionary scenarios. Here we investigated viruses found in parasitic flatworms and their free-living relatives to provide an initial assessment of the viral diversity associated with Platyhelminthes with contrasting lifestyles.

Platyhelminthes, also known as flatworms, constitute a diverse phylum estimated to contain up to 100,000 species with diverse body-plans, life-style and ecological roles ^4,5^. The majority of Platyhelminthes are classified within the Rhabditophora subphylum. Ancestral members of this subphylum have a free-living life-style, including tricladida species used for cellular biology research investigating stem cell, aging, tissue regeneration and homeostasis ^6–8^ (Figure 1). On the other hand, the superclass Neodermata, representing more than half of Platyhelminthes biodiversity, groups endoparasitic trematodes (Digenea and Aspidogastrea) and tapeworms (Cestoda), and ectoparasitic monogeneans (Polyopisthocotylea and Monopisthocotylea)^9^. Here we refer to flatworms outside of Neodermata as ‘free-living’ to distinguish this group with strict parasitic lifestyles. However, note that there are some ‘non-free-living’ flatworms outside of Neodermata.

**Figure 1:**
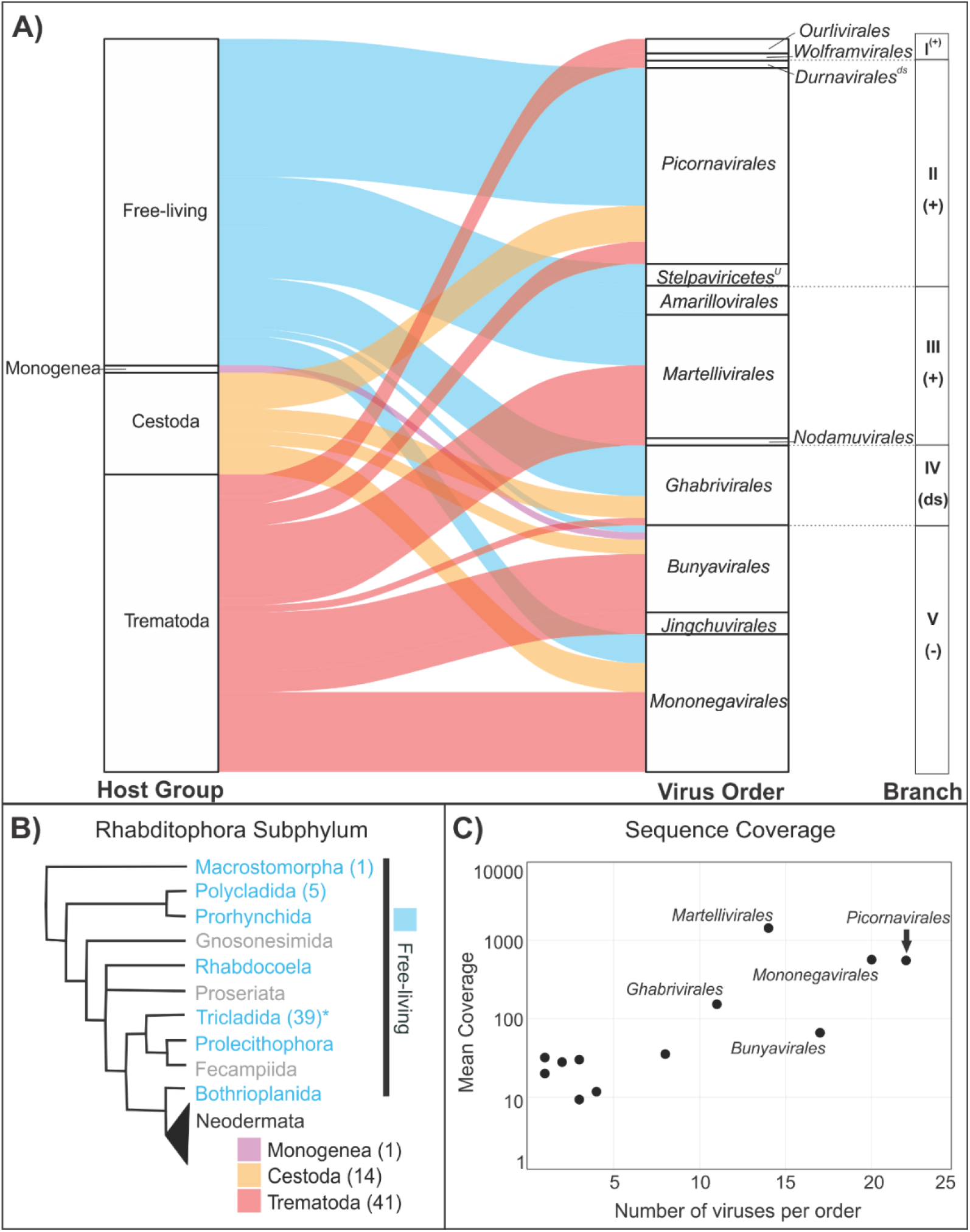
Diversity of viruses discovered in transcriptomic data from 31 species of Platyhelminthes. A) Alluvial plot depicting the distribution of viral sequences identified within flatworm host groups. Viral sequences representing each of the five major branches of the RNA virosphere were grouped at the order level, with the exception of an unassigned (^U^) order within the class *Stepalviricetes*. Branches include members of the *Lenarviricota* (I), *Pisuviricota* (II), *Kitrinoviricota* (III), *Duplornaviricota* (IV), and *Negarnaviricota* (IV) phyla and include positive (+) and (-) sense single-stranded RNA and double-stranded (ds) RNA viruses. Note that *Durnavirales* is the only order within *Pisuviricota* composed of dsRNA viruses. B) Schematic cladogram showing relationships among groups of the Rhabditophora subphylum according to^45^. Cladogram colors correspond to the alluvial plot, where groups highlighted in blue font represent free-living Rhabditophora host groups investigated here (with the exception of Tricladida which includes one species, *Bdelloura candida*, considered to be an ectocommensal* ^46^. Pink, yellow, and red colors represent strict parasitic groups within Neodermata. Rhapditophora taxa in grey font were not investigated here. Numbers within parenthesis indicate the number of viruses detected in a given taxa. C) Mean coverage for viral sequences identified here summarized at the order level.

Neodermatan parasites have contrasting live cycles and are economically relevant pathogens. While monogeneans typically have a single vertebrate fish host, trematodes and tapeworms have complex life-cycles that involve a range of vertebrates as definitive hosts, aninvertebrate intermediate host (typically mollusks and crustaceans, but also cnidarians or polychaetes) and, sometimes, a second intermediate host (mollusks, crustaceans, plants, fish, amphibians) ^9–12^. Neodermatans represent a significant economical and health burden given that these parasites infect humans, cattle and other domesticated animals. For instance, Schistosomiasis, caused by several species of trematodes of the genus *Schistosoma* is the second most important neglected tropical disease after malaria, affecting over 200 million people and causing 200,000 death annually worldwide^13^. Chronic opisthorchiasis and clonorchiasis caused by the liver flukes *Opisthorchis viverrini, Opisthorchis felineus* and *Clonorchis siniensis* have been classified as Group I carcinogens by the International Agency for Research on Cancer due to the increased risk of cholangiocarcinoma associated with infection^14,15^. Humans are also subject to infection by several species of tapeworms including *Taenia spp., Echinococcus spp*., and *Diphyllobothrium spp* ^13,16,17^.

Although our knowledge of flatworm-associated viruses remains very limited, there is evidence indicating that flatworms harbor a diversity of viruses. The very first report of virus-like particle in parasitic Platyhelminthes date from 1976, with observation of geometric arrangements of viral particle in parenchyme cells ^18^. Since then, a few more studies have reported the presence of virus-like particles in monogenean parasites^19,20^ and other flatworms^21–23^. The rise of next generation sequencing technologies has led to the discovery and complete genomic characterization of a ssDNA virus ^24^, a large nidovirus ^25^, and a new family of toti-like viruses^26^ in free-living flatworms. Viruses of the order *Bunyavirales* and the family *Nyamiviridae* (order *Mononegavirales*) have been reported from *Schistosoma japonicum* and a mix of *Taenia sp*. ^27^. More recently, a comprehensive study investigating the virome of the cestode *Schistocephalus solidus* demonstrated that parasitic flatworms may be associated with a large diversity of viruses ^28^. This single species was shown to host multiple species of Rhabdovirus, Nyamivirus, Jingchuvirus, Bunya-like virus, and toti-like viruses^28^.

Herein, we screened for the presence of RNA viruses in the transcriptomes of a broad range of flatworm species. By comparing the phylogenetic position of viruses discovered in ancestral free-living Platyhelminthes and in Neodermatan parasites, we explore the impact of the transition to parasitism on Platyhelminthes virome composition. In addition, we investigate the role of parasite ecology and evolution in virus evolution. When closely related viruses were found, we investigated whether viruses codiversify with their parasitic hosts. Neodermatans could provide opportunities for viruses to complete major host shifts across distantly related taxa given that these parasitic flatworms infect different hosts over the course of their life cycle. Accordingly, we discuss whether parasite viruses have spillover potential based on their evolutionary history.

## Methods

### Building a library of Platyhelminthes transcriptomes

To discover viruses of Platyhelminthes, we downloaded from the Transcriptome Shotgun Assembly (TSA) sequence database all 45 assembled transcriptomes available, corresponding to 38 flatworm species, (Supplementary Table 1). In addition, we downloaded 104 Sequence Read Archive (SRA) files (Supplementary Table 1) from trematodes and cestodes for which no assembled transcriptome was available, which allowed us to process data for an additional 28 species. All SRA datasets were assembled in-house. For this purpose, raw sequences were trimmed for quality and adapter removal using Trimmomatic version 0.36.0 ^29^ with default parameters. Sequence quality after trimming was verified with FastQC version 0.11.5 ^30^ and sequences were assembled with rnaSPAdes as implemented in the SPAdes assembler version 3.11.1 ^31^.

### Searching for viruses in Platyhelminthes transcriptomes

Initially, viral contigs larger than 500 bp were discovered in TSA and assembled SRA data by comparing (BLASTx, e-value < 10^-10^) against a viral protein database containing sequences from the NCBI Reference Sequence database (RefSeq Release number 93, https://www.ncbi.nlm.nih.gov/refseq/). All putative viral transcripts were then translated into proteins to conduct reciprocal BLASTp against the non-redundant (nr) nucleotide and protein database and confirm virus discovery (Supplementary Table 2). When partial viral genomes were identified in TSA datasets, corresponding raw reads were downloaded from SRA and re-assembled in-house as described above. Next, viral contigs within in-house re-assembled transcriptomes were identified through BLASTx against a viral protein database containing flatworm-associated viral sequences initially discovered from TSA and SRA data and closely related sequences. Re-assembly efforts and BLAST searches allowed us to complete or extend the length of partial genome sequences by obtaining longer contigs from in-house assemblies and/or identifying smaller contigs that were used for scaffolding. Viral genome structure and sequence was validated and eventually corrected by mapping all reads and assembled contigs against the newly identified viral genomes as references using BWA (version 0.7.8) ^32^, and visualization using Integrative Genomics Viewer (IGV)^33,34^. When a given genome structure appeared imperfect, an additional assembly, complementary to the initial SPAdes assembly, was obtained with MIRA^35^. Rounds of manual genome sequence correction, read alignment and visualization were conducted. The complete list of viral sequences identified within this study is provided in Supplementary Table 3. Final genome annotations were obtained through BLASTx against RefSeq Release number 207. The distribution of final viral sequences among investigated flatworm hosts was visualized using an alluvial plot prepared in R version 4.0.5. (https://cran.r-proiect.org/web/packages/ggalluvial/vignettes/ggalluvial.html).

### Virus genome characterization and phylogenetic analyses

Open reading frame (ORF) predictions were obtained using Translate on ExPASy and from alignments with related reference virus genomes. Annotation of domains was extracted from comparisons against the Conserved Domain Database (CDD) as implemented by BLASTp against the nr protein database. Initial supergroup assignation was determined from best BLAST matches. Viral RNA-dependent RNA polymerase (RdRP) sequences were aligned using the E-INS-I algorithm implemented in the program MAFFT (version 7) ^36^ to representative sequences of all viral families and genera ratified by the ICTV, as well as additional newly described taxa from recent metatranscriptomic studies ^37–40^. Ambiguously aligned regions were removed using TrimAl (version 1.2) ^41^. For each dataset, the best-fit model of amino acid substitution was determined using Smart Model Selection (SMS) as implemented in PhyML(version 3.0) ^42^. Phylogenetic trees were then inferred using the maximum likelihood method implemented in PhyML (version 3.0) ^42^ using the best-fit model and best of NNI and Subtree Pruning and Regrafting (SPR) branch swapping. Support for nodes on the trees were assessed using an approximate likelihood ratio test (aLRT) with the Shimodaira-Hasegawa-like procedure. Viruses were tentatively taxonomically classified whenever possible based on their phylogenetic position, pairwise sequence identities, pairwise sequence comparison^43^ and/or species demarcation thresholds set by the ICTV (Supplementary Table 3).

### Searching for parasite transcripts in the source data of suspected parasite viruses

Our phylogenetic analysis revealed that some viruses associated with non-parasitic hosts were very closely related to viruses of Neodermatan parasites. To determine whether the detection of these previously reported viral sequences resulted from contamination due to parasitized hosts at the time of sampling, we investigated the presence of parasite sequences. To do this, SRA datasets SRR6291374, SRR6291293, SRR6291349, SRR6291357, and SRX1712840 were downloaded and assembled as described above using SPAdes. Transcript annotation was conducted using MegaBLAST against GenBank to identify transcripts of Neodermatans. In addition, taxonomic composition from each dataset was assessed by comparing 1% of the reads against GenBank. The taxonomic composition was visualized using Krona chart ^44^. The presence of Neodermatans was confirmed when: 1) reads aligned against a parasite whose ecology matched with the sample and 2) transcripts had significant BLAST matches to Neodermatan parasite proteins (i.e. cytochrome c oxidase, transcription elongation factor, HSP70, ATPase)

## Results and Discussion

### Platyhelminthes harbor a diverse RNA virome

We conducted a large-scale survey of Platyhelminthes-associated viruses through data mining of TSA and SRA publicly available transcriptomes. In total, 149 datasets, corresponding to 66 flatworm species representing free-living Rhabditophora and parasitic Neodermata, were screened for viruses (Supplementary Table 1). Viruses were successfully detected within datasets from free-living Rhabditophora (45 viruses) and Nematoda, including Trematoda (41 viruses), Cestoda (14 viruses) and Monogenea (1 virus) (Figure 1). A total of 115 unique sequences with either complete (87 sequences) or partial (28 sequences) protein coding sequence regions were identified, representing 101 novel viruses, with a small minority of viruses with fragmented genomes (Supplementary Tables 2 and 3). Importantly, the investigated viral sequences were not found in available Platyhelminthes genomes and encoded for proteins without frameshifts, nonsense mutations, or repeat sequences that are common in endogenous viral elements. Therefore, viruses described here are most likely exogenous functional viruses.

The novel Platyhelminthes-associated virus species were distributed among all five major Phyla of RNA viruses, and fell within a total of twelve orders revealing the large diversity of virus taxa found within flatworms (Figure 1, Supplementary table 4). Only two viruses were classified at the genus level (Nyamiviridae family, genus *Tapwovirus)* and 34 viruses at the family level, indicating that Platyhelminthes host a unique viral diversity. The majority of viruses were classified at the order level, including *Picornavirales* (27 viruses), *Mononegavirales* (21 viruses), *Bunyavirales* (17 viruses)*, Martellivirales* (14 viruses), *Ghabrivirales* (11 viruses), and *Amarillovirales* (8 viruses) (Figure 1). Other orders were represented by five members or less, including *Wolframvirales*, *Ourlivirales*, *Durnavirale*s, an unassigned order of *Stelpaviricetes*, *Amarillovirales*, *Nodamuvirales*, and *Jingchuvirales*. We observed a significant relationship between the number of viruses discovered within an order and the mean read coverage of those viruses (Figure 1C). It is possible that viral taxa with low mean coverage are less represented in flatworms. Alternatively, it is possible that methodological and sequencing approaches used in transcriptomic studies investigated here did not recover viruses with low abundance, which would suggest that Platyhelminthes host an even greater diversity of viruses than presented herein.

We investigated phylogenetic relationships among Platyhelminthes-associated viral taxa. Viruses within unassigned families of the order *Picornavirales, Martellivirales, Ghabrivirales, Bunyavirales* and *Jingchuvirales*, as well as viruses within the families *Flaviviridae, Nyamiviridae* and *Rhabdoviridae* were found within more than one Platyhelminthes species. We used phylogenetic methods to further investigate their relationship to each other, and to the known viral diversity. To build phylogenetic trees, we included representatives of previously characterized families as well as unassigned viruses that showed high sequence similarity to viruses described here. Phylogenetic analyses revealed that viruses of Platyhelminthes often cluster together, and separately from other known viruses, providing evidence that they constitute distinct taxa. Note that newly discovered flatworm-associated viruses that were not phylogenetically related to other viruses of Platyhelminthes were not investigated further because the host remains putative. These viruses could be associated with the host diet, with a co-infecting microorganism, or could result from contamination during sample processing. Overall, including only taxa for which viruses were found in at least two different Platyhelminthes species, our data provide strong evidence for at least seven new families and eighteen new genera of Platyhelminthes specific viruses (Supplementary Figures 1 – 16, Supplementary Table 3, Figure 1).

### Previously reported viruses of Neodermatan parasites within vertebrate and invertebrate hosts

In some instances, the newly identified Neodermatan-associated viruses clustered closely with vertebrate- and invertebrate-associated viruses previously discovered through metagenomic and metatranscriptomic studies^2,27,47^. However, upon close inspection, we found transcripts that belong to Platyhelminthes within the original datasets from these reports indicating that parasites were present at the time of sampling. Specifically, we found transcripts from an unknown trematode (likely from the family Fascioliidae or Dicrocoeliidae) in the dataset from a razor shell specimen (SRR1712840) that contained a picorna-like virus (Beihai razor-shell virus 4) and a bunya-like virus (Beihai bunya-like virus 2) closely related to trematode-associated viruses discovered here. The spotted paddle-tail newt (SRR6291293) within which a Neodermatan-like rhabdovirus was found (Fujian dimarhabdovirus) appeared infected by trematodes known to infect amphibians and non-fish vertebrates (*Mesocelium sp*. and *Spriometra sp.)*. The Wenling sharpspine skate mix (SRR6291349) within which another Neodermatan-like rhabdovirus was found (Wenling dimarhabdovirus 8) was infected by a cestode from the family *Echinobothriidae* known to infect Elasmobranch. Finally, two sample mixes of fish gills (SRR6291357 and SRR6291374) within which more Neodermatan virus-like rhabdoviruses were found (Wenling dimarhabdovirus 10 and Beihai dimarhabdovirus 1), contained reads that align against Monogenean parasite nucleotide sequences, but the low percentage of reads belonging to Platyhelminthes prevented the successful assembly of transcripts. The phylogenetic positions of these viruses, and the demonstration that the host organisms were infected by Neodermatan parasites at the time of sampling strongly suggest that these viruses were actually infecting the parasite rather than the vertebrate or invertebrate host. Additional viruses that were probably associated with tapeworms or fluke but for which we could not conduct the same analysis (either the raw data were not available or samples were processed in a way that eliminated host-associated transcripts) include Picornaviruses (Fesavirus 3^48^, Pernambuco virus^49^, Arivirus 2^50^, Blackbird arilivirus^51^), and an additional rhabdovirus (Fox fecal rhabdovirus^52^). Future studies investigating the virome of vertebrates, mollusks or crustaceans that are either definitive or intermediate host of Neodermatan parasites, need to consider the presence of such stowaway passengers when assigning hosts to newly discovered viruses.

### Neodermatan viruses are distinct from viruses of free-living Rhabditophora

Differences between free-living and parasitic Platyhelminthes were reflected in the type of viruses they harbored. We discovered a greater diversity of positive-strand RNA viruses and double-stranded RNA viruses of the families *Picornavirales* and *Ghabrivirales* in the free-living Rhabditophora than in the parasitic Neodermatans (Figure 1). A similar trend was observed in the only non-free-living flatworm included in the free-living Rhabditophora dataset, an ectocommensal named *Bdelloura candida*. In contrast, Neodermatan parasites harbored a greater diversity of negative-strand RNA viruses. For example, negative-strand RNA viruses of a novel family within the order *Jingchuvirales*, and viruses of the family *Rhabdoviridae*, order *Mononegavirales* were found exclusively in Neodermatan parasites (Figure 1). When viruses of a given clade were found in both ancestral Platyhelminthes and Neodermatan parasites, they clustered separately on phylogenetic trees. This distinct clustering was evident for unassigned viral families of the orders *Picornavirales*, *Bunyavirales* and *Ghabrivirales* (Figure 2). There were only two cases where viruses of Neodermata and Rhabditophora clustered together on the phylogenetic tree. Within the family *Nyamiviridae*, viruses of tapeworms clustered closely together within the genus *Tapwovirus*, whereas viruses of Rhabditophora were more closely related to viruses of the genus *Berhavirus* (Figure 2). Within the order *Martellivirales*, the Psilosi virus found in the trematode *P. similimum* clustered closely with viruses of Rhabditophora (*P. torva* and S. *mediterranea*), whereas the provittati virus of the free-living worm *P. vitatti* clustered most closely with viruses of liver flukes. Yet, the RdRP of these viruses showed a maximum of 36% and 45% aa identity to their closest relatives suggesting that they belong to different taxa. More sampling is needed to help resolve the phylogeny of Platyhelminthes. Nevertheless, our findings indicate that the transition of a proto-neodermatan worm from free-living to parasitism over 500Ma ago ^53^ impacted virus evolution.

**Figure 2:**
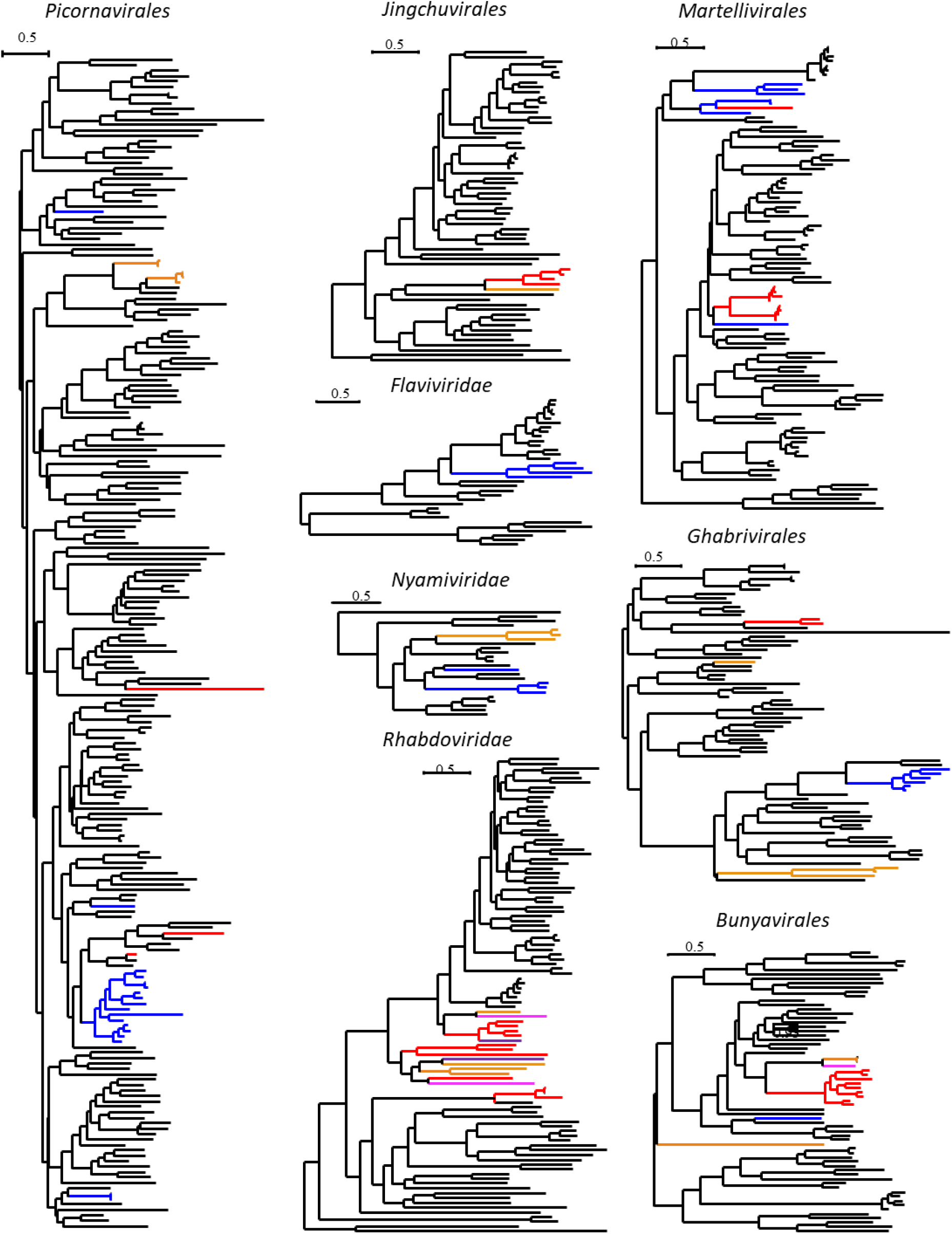
Distinct viruses identified within free-living Rhabditophora and parasitic Neodermata. Phylogenetic trees of the RNA-directed RNA polymerase (RdRPs) of RNA viruses of the orders *Picornavirales, Martellivirales*, *Bunyavirales*, *Ghabrivirales* and *Jingchuvirales*, and the families *Flaviviridae, Nyamiviridae* and *Rhabdovirdae*. Viruses of Platyhelminthes included in the trees are color coded (trematode, red; cestode, orange; monogenean, pink; Rhabditophora, blue). The trees were inferred in PhyML and high-resolution annotated trees are available in Supplementary Figures 1 to 8.

### Viruses of Neodermata co-diversify with their parasitic hosts

Viruses of Neodermata, with the exception of rhabdoviruses (see below), often clustered separately based on their parasitic host’s phylogenetic relationships suggesting a close association between parasitic hosts and their viruses. Within the order Ghabrivirales, viruses of cestodes and trematodes are found on distinct branches and constitute two novel proposed families suggesting distinct evolutionary origins before diversification within their parasitic hosts (Figure 2). Within the order *Jingchuvirales*, Neodermatan viruses showed a maximum of 19% aa identity and clustered separately from all other viruses, suggesting that they constitute a novel family (Figure 2). Among those, the cestode-associated virus Schistocephalus solidus jingchuvirus (SsJV) has only 23-29% aa identity with Jingchuviruses of trematodes indicating that they belong to two distinct genera within the same family. Similarly, viruses of the order *Bunyavirales* associated with trematodes clustered separately (with 19-23% identity) from the viruses associated with the monogenean *Eudiplozoon nipponicum* and the cestode *T. nodulus* (Figure 3). Those two viruses showed a 40% aa identity while Bunya-like viruses of trematodes further clustered depending on the parasite’s family, with 57-71% identity when the host belong to the same family but 37-47% identity when the host belongs to different parasite families. Viruses associated with the Schistosomatidae *(Schistosoma japonicum* and *Trichobillharzia regenti)* clustered together, separately from viruses associated with the liver flukes of the family Opisthorchiidae (*M. orientalis* and *Clonorchis siniensis*) that clustered together, and separately from viruses associated with the Psilostomidae (*Psilotrema similimum* and *S. pseudoglobulus*). Past experimental investigations on *S. solidus* virus transmission mode revealed that most viruses, including SsJV and a Bunya-like virus are vertically transmitted from parents to offspring^54^. The co-diversification of Jingchuviruses and Bunya-like viruses with their cestode and trematode hosts support this finding and indicate that vertical transmission may be common within these new virus taxa.

**Figure 3:**
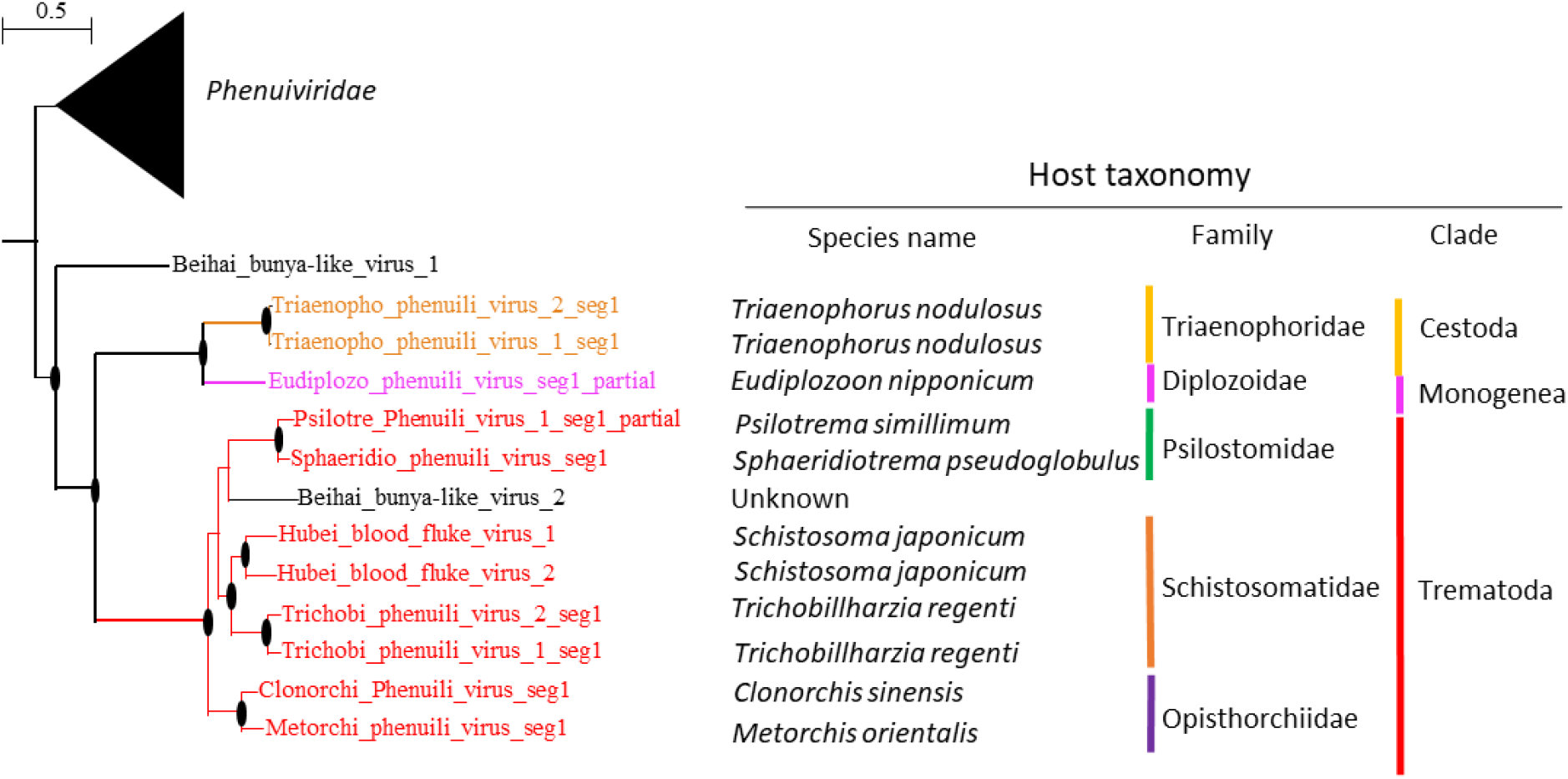
Neodermatan viruses of the order Bunyavirales co-diversify with their parasitic hosts. The figure provides a phylogenetic tree of the RNA-directed RNA polymerase (RdRPs) of RNA viruses of the order Bunyavirales found in Neodermatan parasites and closely related viruses of the family *Phenuiviridae*. Viruses of Platyhelminthes included in the trees are color coded (trematode, red; cestode, orange; Rhabditophora, blue). The tree was inferred in PhyML using the LG substitution model. Branch points indicate that results of Shimodaira-Hasgawa branch test > 0.9. The species name, family and order of the hosts of identified viruses are provided next to the branches.

### Rhabdoviruses exemplify the potential role of parasites in virus evolution

Parasites have intimate relationships with their hosts, and they can infect different host individuals or species over the course of their life-cycle, two factors that can facilitate virus transmission, and spill-over^3^. The phylogenetic position of the diverse novel rhabdoviruses of Neodermatan parasites suggest that these viruses switch host often and can at times emerge in their hosts. Even though Neodermatan-associated rhabdoviruses mostly clustered together, we did not observe a distinct co-diversification with their hosts (Figure 4). Indeed, our analysis suggests that the thirteen novel rhabdoviruses belong to a minimum of seven distinct taxa (Figure 4, Supplementary Figure 8). There was no obvious clustering of viruses of cestodes and trematodes indicating that virus host-shifts occurred on multiple occasions. Hahn et al. ^54^ recently showed that the rhabodvirus SsRV1 is excreted by adult *S. solidus* and is transmitted to parasitized hosts. If this characteristic is conserved among Neodermatan-associated rhabdoviruses, it would provide an avenue for rhabdovirus host switching between parasites co-infecting the same hosts and between parasites and their intermediate and definitive hosts.

**Figure 4:**
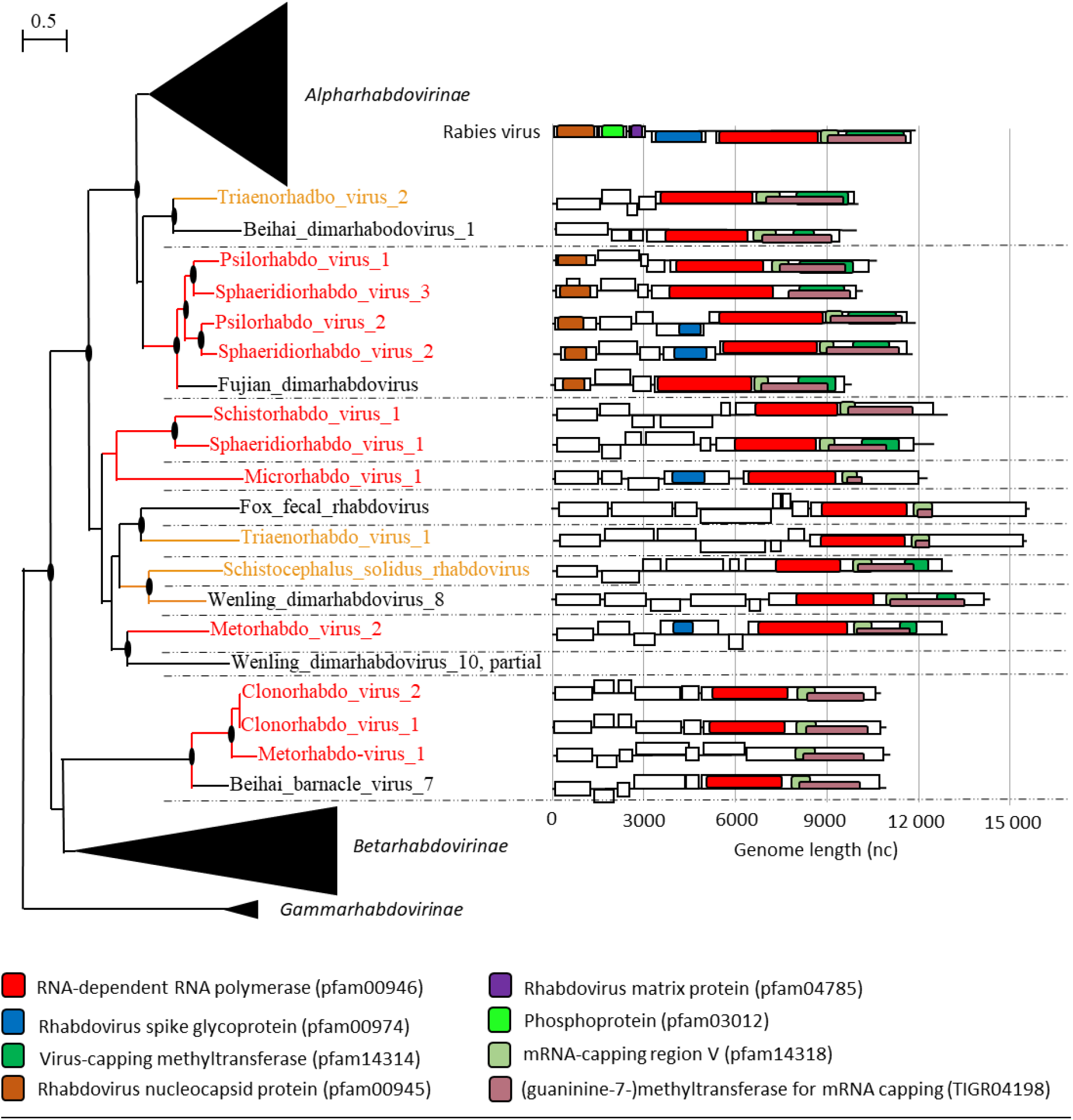
Neodermatan viruses of the family *Rhabdoviridae*. A. Phylogenetic trees of the RNA-directed RNA polymerase (RdRPs) of RNA viruses of the family *Rhabdoviridae*. Viruses of Platyhelminthes included in the trees are color coded (trematode, red; cestode, orange). The dotted lines delineate different taxa. The tree was inferred in PhyML using the LG substitution model. Branch points indicate that results of Shimodaira-Hasgawa branch test > 0.9. B. Genome organization of Neodermatan viruses of the family *Rhabdoviridae*. Boxes represent putative genes. The black line indicates noncoding region.

The genomes of rhabdoviruses of cestodes and trematodes exhibited extensive diversity in genome length (from 9963 nt to 15554 nt). As expected, this genome length variation was associated with variation in the number of predicted genes. The rhabdovirus genome is normally composed of a minimum of five ORFs encoding for five structural proteins – the nucleoprotein (N), the polymerase-associated phosphoprotein (P), the matric protein (M), the glycoprotein (G) and the RdRP (L). Yet, five of the rhabdoviruses of Neodermata encoded only four proteins, with the loss of the spike glycoprotein. Other Neodermata rhabodvirus genomes contained up to three additional small ORFs between G and L encoding putative proteins.

When looking at a broader resolution, all the known diversity of alpharhabdoviruses and betarhabdoviruses is nested within the diversity of viruses of Platyhelminthes, with gammarhabdoviruses of fish maintaining an ancestral position to all rhabdoviruses. This is notable considering that alpharabdoviruses contain vertebrate and vector-borne rhabdoviruses and betarhabdoviruses encompass plant and arthropod viruses. A parsimonious explanation for this phylogenetic distribution would be that an ancestral Neodermatan parasite initially acquired a rhabdovirus from its fish host. Then, over the course of Neodermatan diversification and increasing range of intermediate and definitive hosts, these viruses have switch host again, giving rise to the diversity of alpharhabdoviruses and betarhabdoviruses known to date. Most specifically, our phylogenetic analysis shows that rhabdoviruses of Neodermatan parasites are close ancestors to Lyssaviruses, indicating that a parasite-associated rhabdovirus emerged in a parasitized vertebrate host and became the ancestral Lyssavirus. Clearly, throughout their evolutionary history, rhabdoviruses have maintained the ability to switch host frequently^55^. Host-switching would explain why rhabdoviruses often emerge, or re-emerge, as zoonotic and epizootic viral diseases^56^ and the diversity of host associations within the alpharhabdovirinae subfamily^55^. Our analysis indicates that Neodermatan-associated rhabdoviruses should be considered as a potential source of viral emergence.

### Role of viruses in parasite infection and perspectives

Much remains to be learned about parasitic flatworm viruses and their role in parasite ecology and evolution. Which viruses of Neodermata can infect parasitized vertebrate and invertebrate hosts? Can these viruses be responsible for symptoms, and associated pathologies that have been attributed to the parasitic flatworms? Does viral infection have a deleterious effect on parasite fitness? Alternatively, can viruses increase parasitic flatworm reproduction and transmission? What is the effect of co-infection by a parasite and its associated virus on host immune response? How viruses contribute to host-parasitic flatworm co-evolution?

Here we took advantage of publicly available transcriptomic data to identify viruses. However, transcriptomic data generated from parasitic flatworms, or their hosts, could also be used to gain additional information on virus prevalence, transmission to the host, or cellular location. For instance, we used a series of transcriptomic datasets^57,58^ to investigate a novel dicistro-like virus named Schmimed virus 1. The transcriptomic data revealed Schmimed virus 1 neurotropism and the role of the Hippo pathway in controlling virus replication within its planarian host (Supplementary Figure 17).

Gaining knowledge about Neodermatan viral infections would allow us to understand how viruses may affect parasite-host interactions. Viruses could play a role in parasitic flatworm infections as exemplified by three-partite interactions in other systems. In the parasitoid wasp *Leptopilina boulardii*, the Leptopolina boulardi filamentous virus (LbFV) manipulates the parasitic wasp behavior to increase hyperparasitism - forcing the wasp to lay its eggs in already parasitized hosts – and increase egg load to increase its horizontal transmission^59–61^. Another parasitoid wasp, *Dinocampus coccinellae* transmits a neurotropic RNA viruses to its coccinellid host to manipulate the host behavior and force it to protect the parasite progeny^62^. In *Leishmania (Vianna)*, the Leishmania RNA virus 1 (LRV1) is excreted within exosomes and exacerbates *Leishmania* pathogenicity, by causing a hyper-inflammatory response and metastatic secondary lesions, known as muco-cutaneous leishmaniasis ^63,64^. In another protozoan, *Trichomonas vaginalis*, the Trichomonavirus (TVV)-induced pro-inflammatory innate immune response is amplified upon anti-parasitic treatment due to the release of viruses by dying parasites ^65,66^. Clearly, the potential role of viruses in Neodermatan pathogenicity and in symptoms associated with anti-parasitic treatments is broad and merits in-depth investigation^67–69^. Depending on the nature of virus-parasite interactions and impact on the parasitized host, viruses may be used as biocontrol agents to reduce parasite population size, or they may be targets of new vaccines or antiviral treatments to reduce parasite pathogenicity. Characterizing viruses of parasites also offer new opportunities in functional genomics. It has been proposed that viruses of parasites could be used to produce pseudotyped viruses with high specificity to the target parasite species to produce stable lines of transgenic parasites^70^.

## Authors contribution

NMD conceived and designed the study, acquired data, contributed to data analysis and interpretation, prepared figures, and wrote the first draft of the manuscript. KR acquired data, contributed to data analysis and interpretation, edited manuscript, and prepared figures. YB contributed to data analysis and edited manuscript. PL contributed to data analysis. All authors revised the manuscript and approved the final version.

## Acknowledgment

This project was part of the Parasite Microbiome project. NMD was supported by project SAD19002 SANIBIOM provided by Region Bretagne, Direction du Développement Economique.

## Data availability

Viral sequences have been submitted to NCBI and accession numbers will be provided upon publication.

