## Supplementary figures 1-17 for "A world of viruses nested within parasites: Unraveling viral diversity within parasitic flatworms (Platyhelminthes)"

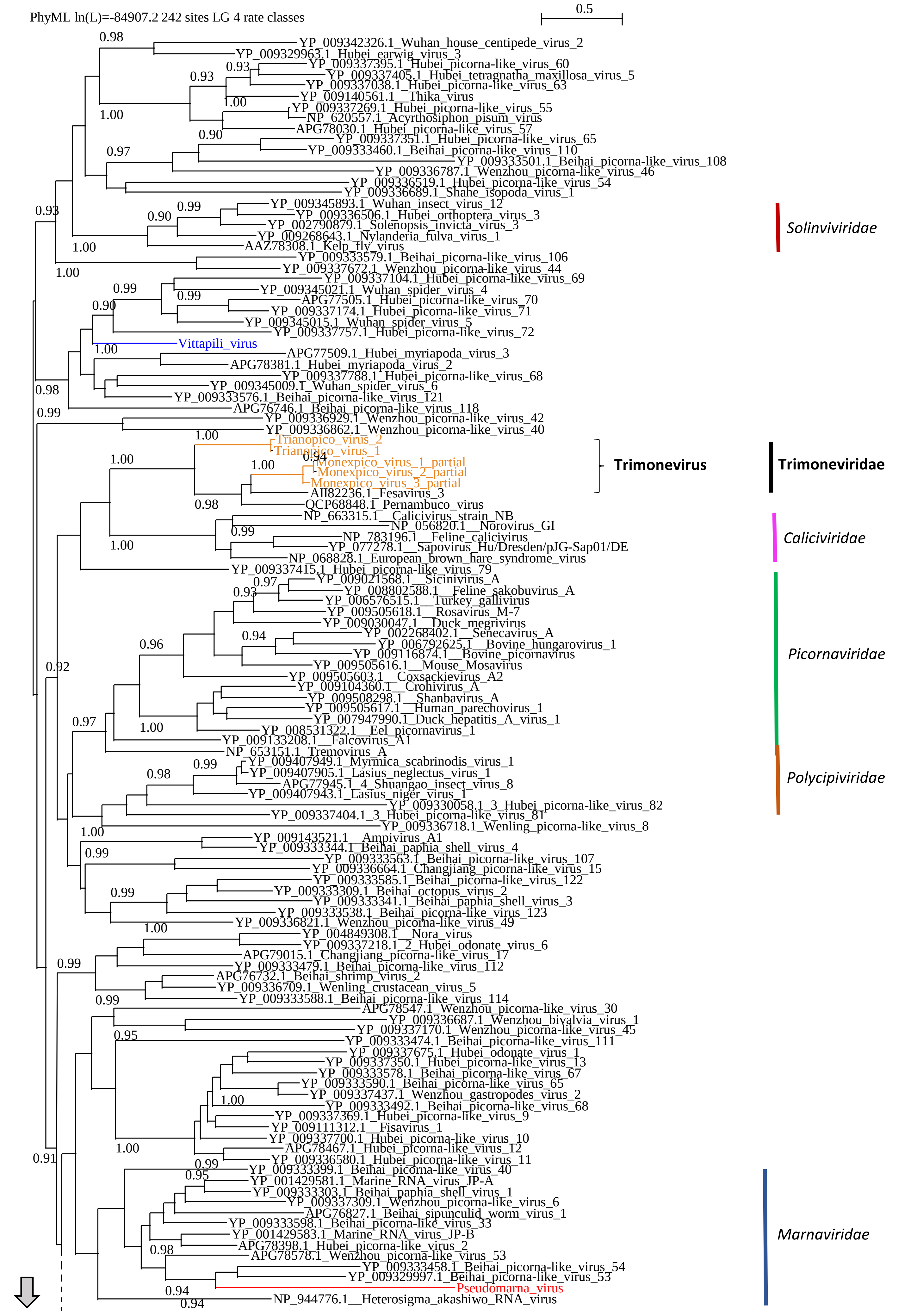
PhyML ln(L)=-84907.2 242 sites LG 4 rate classes
0.5
YP_009342326.1_Wuhan_house_centipede_virus_2
YP_009329963.1_Hubei_earwig_virus_3
YP_009337395.1_Hubei_picorna-like_virus_60
YP_009337405.1_Hubei_tetragnatha_maxillosa_virus_5
YP_009337038.1_Hubei_picorna-like_virus_63
YP_009140561.1__Thika_virus
YP_009337269.1_Hubei_picorna-like_virus_55
NP_620557.1_Acyrthosiphon_pisum_virus
APG78030.1_Hubei_picorna-like_virus_57
YP_009337351.1_Hubei_picorna-like_virus_65
YP_009333460.1_Beihai_picorna-like_virus_110
YP_009333501.1_Beihai_picorna-like_virus_108
YP_009336787.1_Wenzhou_picorna-like_virus_46
YP_009336519.1_Hubei_picorna-like_virus_54
YP_009336689.1_Shahe_isopoda_virus_1
YP_009345893.1_Wuhan_insect_virus_12
YP_009336506.1_Hubei_orthoptera_virus_3
YP_002790879.1_Solenopsis_invicta_virus_3
YP_009268643.1_Nylanderia_fulva_virus_1
AAZ78308.1_Kelp_fly_virus
Solinviviridae
YP_009333579.1_Beihai_picorna-like_virus_106
YP_009337672.1_Wenzhou_picorna-like_virus_44
YP_009337104.1_Hubei_picorna-like_virus_69
YP_009345021.1_Wuhan_spider_virus_4
APG77505.1_Hubei_picorna-like_virus_70
YP_009337174.1_Hubei_picorna-like_virus_71
YP_009345015.1_Wuhan_spider_virus_5
YP_009337757.1_Hubei_picorna-like_virus_72
Vittapili_virus
APG77509.1_Hubei_myriapoda_virus_3
APG78381.1_Hubei_myriapoda_virus_2
YP_009337788.1_Hubei_picorna-like_virus_68
YP_009345009.1_Wuhan_spider_virus_6
YP_009333576.1_Beihai_picorna-like_virus_121
APG76746.1_Beihai_picorna-like_virus_118
YP_009336929.1_Wenzhou_picorna-like_virus_42
YP_009336862.1_Wenzhou_picorna-like_virus_40
Trianopico_virus_2
Trianopico_virus_1
Monexpico_virus_1_partial
Monexpico_virus_2_partial
Monexpico_virus_3_partial
AII82236.1_Fesavirus_3
QCP68848.1_Pernambuco_virus
Trimonevirus
Trimoneviridae
NP_663315.1__Calicivirus_strain_NB
NP_056820.1__Norovirus_GI
NP_783196.1__Feline_calicivirus
YP_077278.1__Sapovirus_Hu/Dresden/pJG-Sap01/DE
NP_068828.1_European_brown_hare_syndrome_virus
Caliciviridae
YP_009337415.1_Hubei_picorna-like_virus_79
YP_009021568.1__Sicinivirus_A
YP_008802588.1__Feline_sakobuvirus_A
YP_006576515.1__Turkey_gallivirus
YP_009505618.1__Rosavirus_M-7
YP_009030047.1__Duck_megrivirus
YP_002268402.1__Senecavirus_A
YP_006792625.1__Bovine_hungarovirus_1
YP_009116874.1__Bovine_picornavirus
YP_009505616.1__Mouse_Mosavirus
YP_009505603.1__Coxsackievirus_A2
YP_009104360.1__Crohivirus_A
YP_009508298.1__Shanbavirus_A
YP_009505617.1__Human_parechovirus_1
YP_007947990.1_Duck_hepatitis_A_virus_1
YP_008531322.1__Eel_picornavirus_1
YP_009133208.1__Falcovirus_A1
NP_653151.1_Tremovirus_A
Picornaviridae
YP_009407949.1_Myrmica_scabrinodis_virus_1
YP_009407905.1_Lasius_neglectus_virus_1
APG77945.1_4_Shuangao_insect_virus_8
YP_009407943.1_Lasius_niger_virus_1
YP_009330058.1_3_Hubei_picorna-like_virus_82
YP_009337404.1_3_Hubei_picorna-like_virus_81
YP_009336718.1_Wenling_picorna-like_virus_8
Polycipiviridae
YP_009143521.1__Ampivirus_A1
YP_009333344.1_Beihai_paphia_shell_virus_4
YP_009333563.1_Beihai_picorna-like_virus_107
YP_009336664.1_Changjiang_picorna-like_virus_15
YP_009333585.1_Beihai_picorna-like_virus_122
YP_009333309.1_Beihai_octopus_virus_2
YP_009333341.1_Beihai_paphia_shell_virus_3
YP_009333538.1_Beihai_picorna-like_virus_123
YP_009336821.1_Wenzhou_picorna-like_virus_49
YP_004849308.1__Nora_virus
YP_009337218.1_2_Hubei_odonate_virus_6
APG79015.1_Changjiang_picorna-like_virus_17
YP_009333479.1_Beihai_picorna-like_virus_112
APG76732.1_Beihai_shrimp_virus_2
YP_009336709.1_Wenling_crustacean_virus_5
YP_009333588.1_Beihai_picorna-like_virus_114
APG78547.1_Wenzhou_picorna-like_virus_30
YP_009336687.1_Wenzhou_bivalvia_virus_1
YP_009337170.1_Wenzhou_picorna-like_virus_45
YP_009333474.1_Beihai_picorna-like_virus_111
YP_009337675.1_Hubei_odonate_virus_1
YP_009337350.1_Hubei_picorna-like_virus_13
YP_009333578.1_Beihai_picorna-like_virus_67
YP_009333590.1_Beihai_picorna-like_virus_65
YP_009337437.1_Wenzhou_gastropodes_virus_2
YP_009333492.1_Beihai_picorna-like_virus_68
YP_009337369.1_Hubei_picorna-like_virus_9
YP_009111312.1__Fisavirus_1
YP_009337700.1_Hubei_picorna-like_virus_10
APG78467.1_Hubei_picorna-like_virus_12
YP_009336580.1_Hubei_picorna-like_virus_11
YP_009333399.1_Beihai_picorna-like_virus_40
YP_001429581.1_Marine_RNA_virus_JP-A
YP_009333303.1_Beihai_paphia_shell_virus_1
YP_009337309.1_Wenzhou_picorna-like_virus_6
APG76827.1_Beihai_sipunculid_worm_virus_1
YP_009333598.1_Beihai_picorna-like_virus_33
YP_001429583.1_Marine_RNA_virus_JP-B
APG78398.1_Hubei_picorna-like_virus_2
APG78578.1_Wenzhou_picorna-like_virus_53
YP_009333458.1_Beihai_picorna-like_virus_54
YP_009329997.1_Beihai_picorna-like_virus_53
Pseudomarna_virus
NP_944776.1__Heterosigma_akashiwo_RNA_virus
Marnaviridae

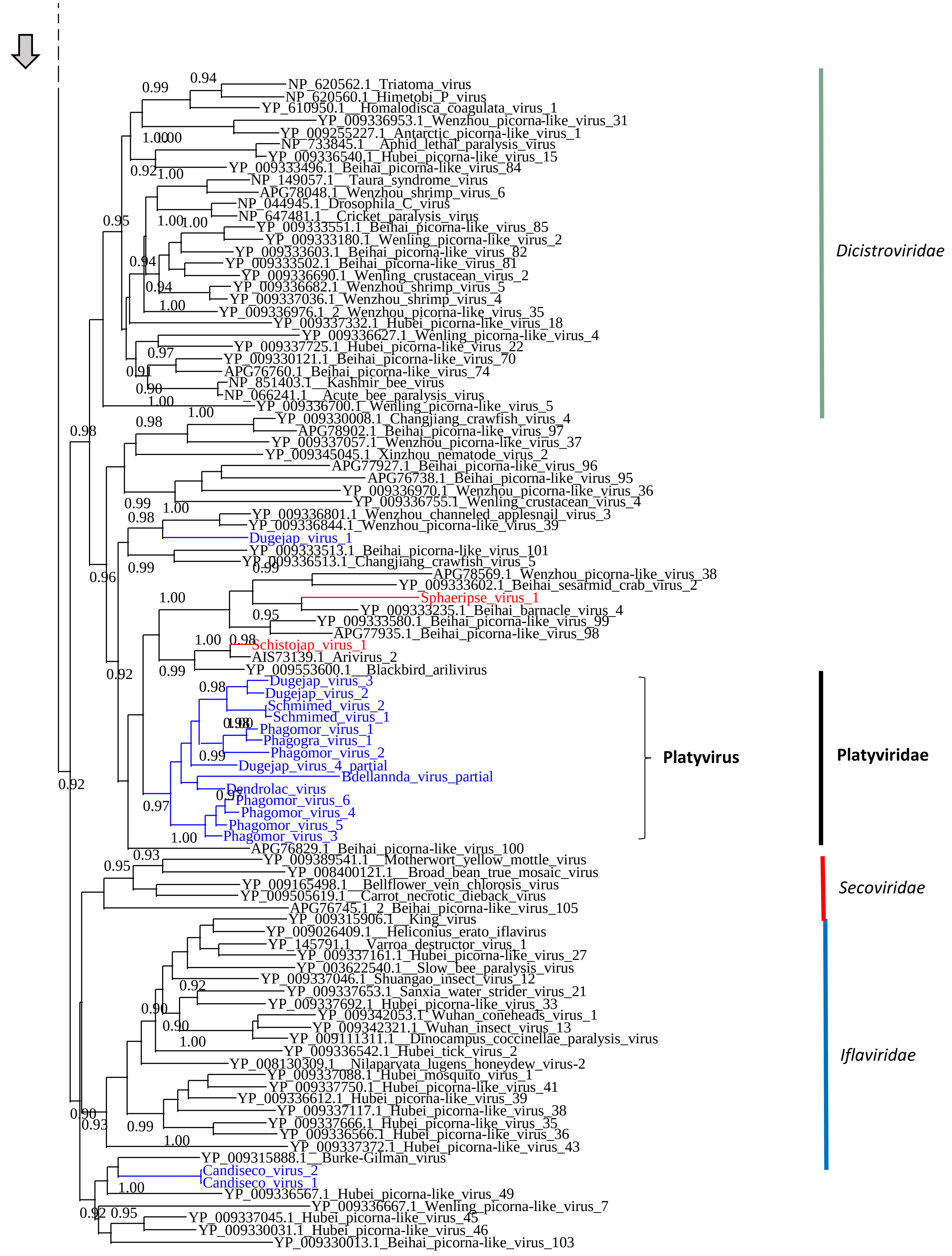


Supplementary Figure 1: Phylogenetic trees of the RNA-directed RNA polymerase (RdRPs) of RNA viruses of the order *Picornavirales.* Viruses of Platyhelminthes included in the trees are color coded (trematode, red; cestode, orange; monogenean, pink; Rhabditophora, blue). The trees were inferred in PhyML using the LG substitution model. Branch points indicate that results of Shimodaira-Hasgawa branch test > 0.9.

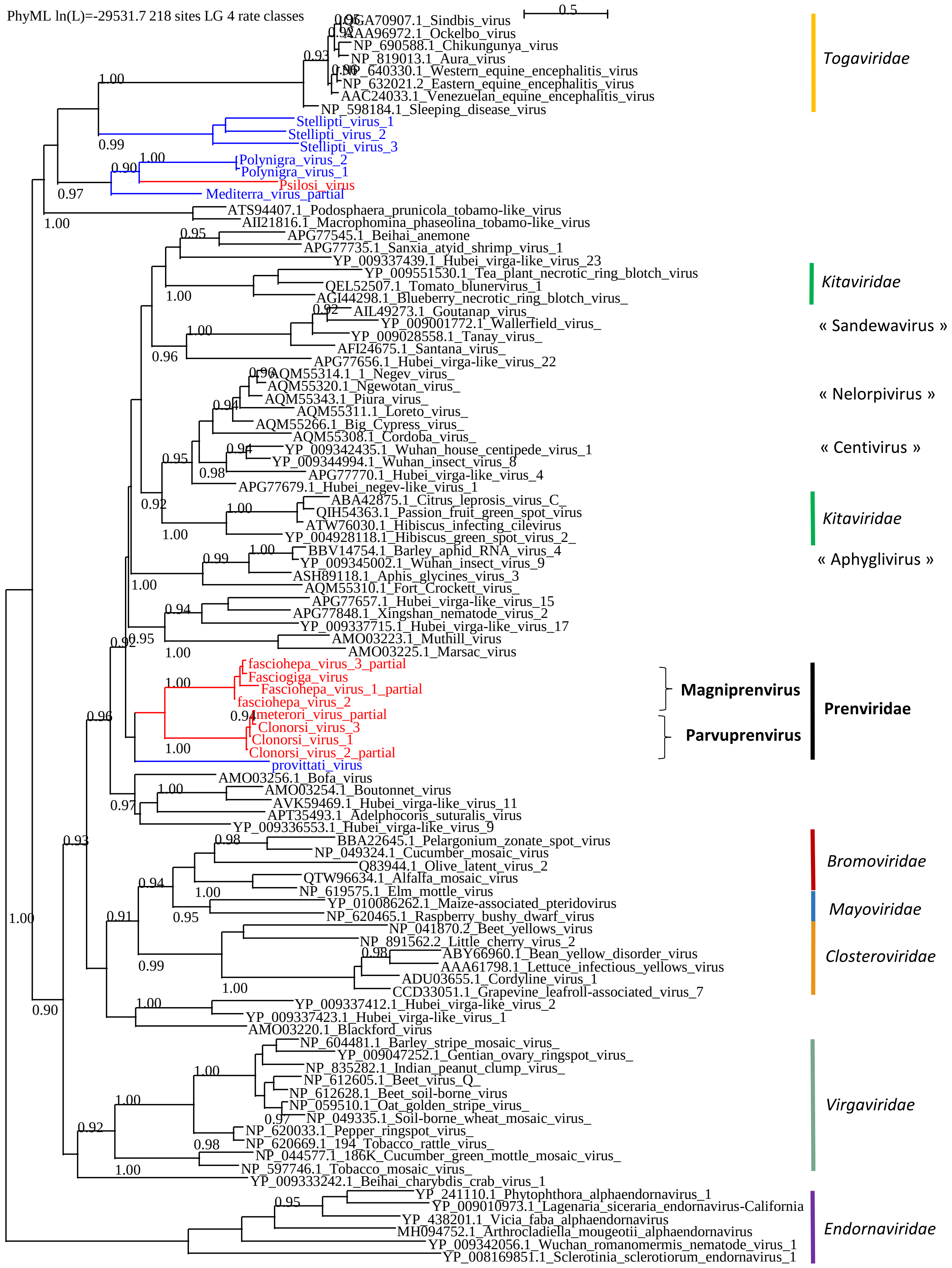


Supplementary Figure 2: Phylogenetic trees of the RNA-directed RNA polymerase (RdRPs) of RNA viruses of the order *Martellivirales.* Viruses of Platyhelminthes included in the trees are color coded (trematode, red; cestode, orange; monogenean, pink; Rhabditophora, blue). The trees were inferred in PhyML using the LG substitution model. Branch points indicate that results of Shimodaira-Hasgawa branch test > 0.9. Assigned family names are provided in Italic. Novel proposed genus and family names are indicated in bold and black

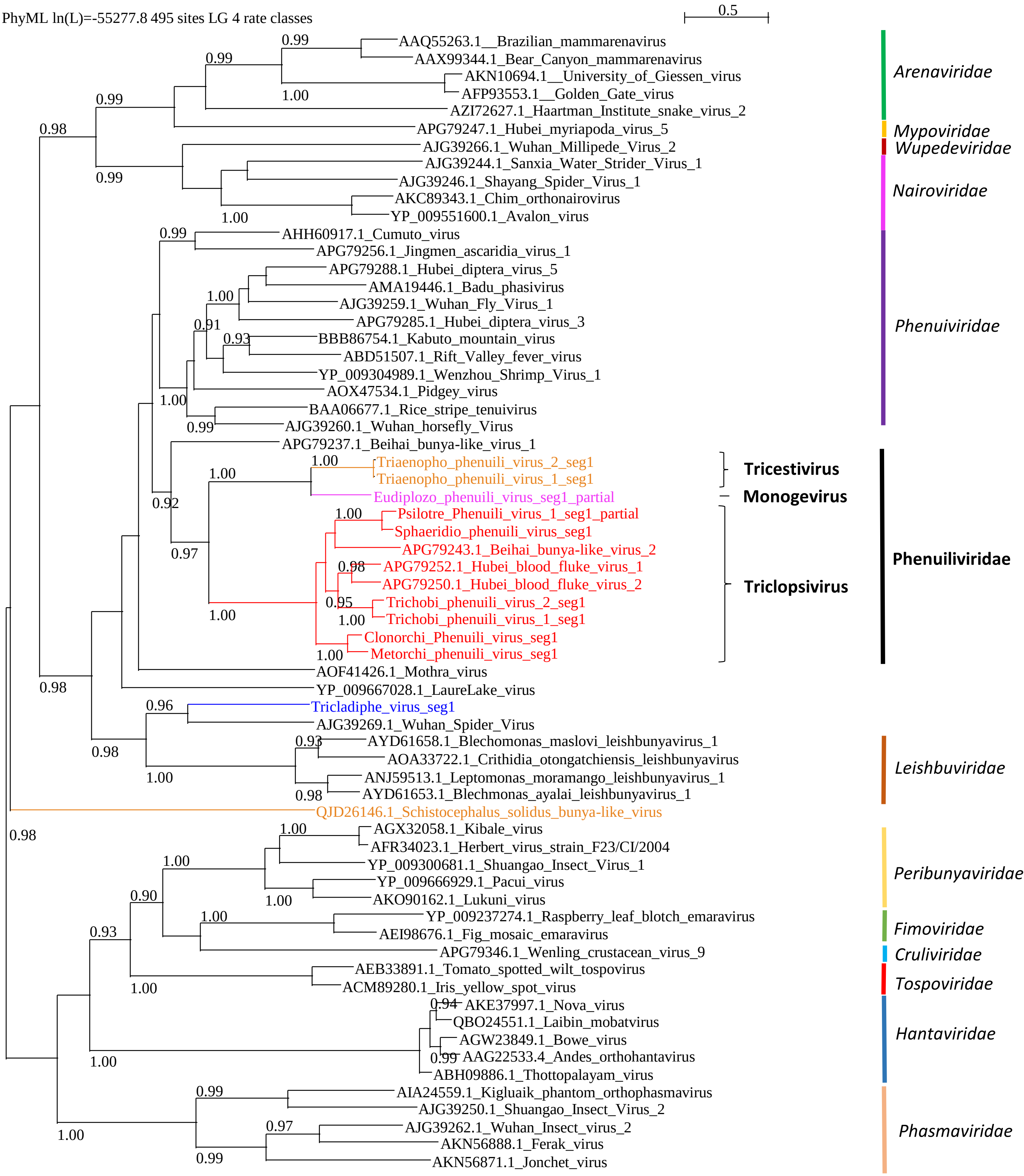


Supplementary Figure 3: Phylogenetic trees of the RNA-directed RNA polymerase (RdRPs) of RNA viruses of the order *Bunyavirales.* Viruses of Platyhelminthes included in the trees are color coded (trematode, red; cestode, orange; monogenean, pink; Rhabditophora, blue). The trees were inferred in PhyML using the LG substitution model. Branch points indicate that results of Shimodaira-Hasgawa branch test > 0.9. Assigned family names are provided in Italic. Novel proposed genus and family names are indicated in bold and black

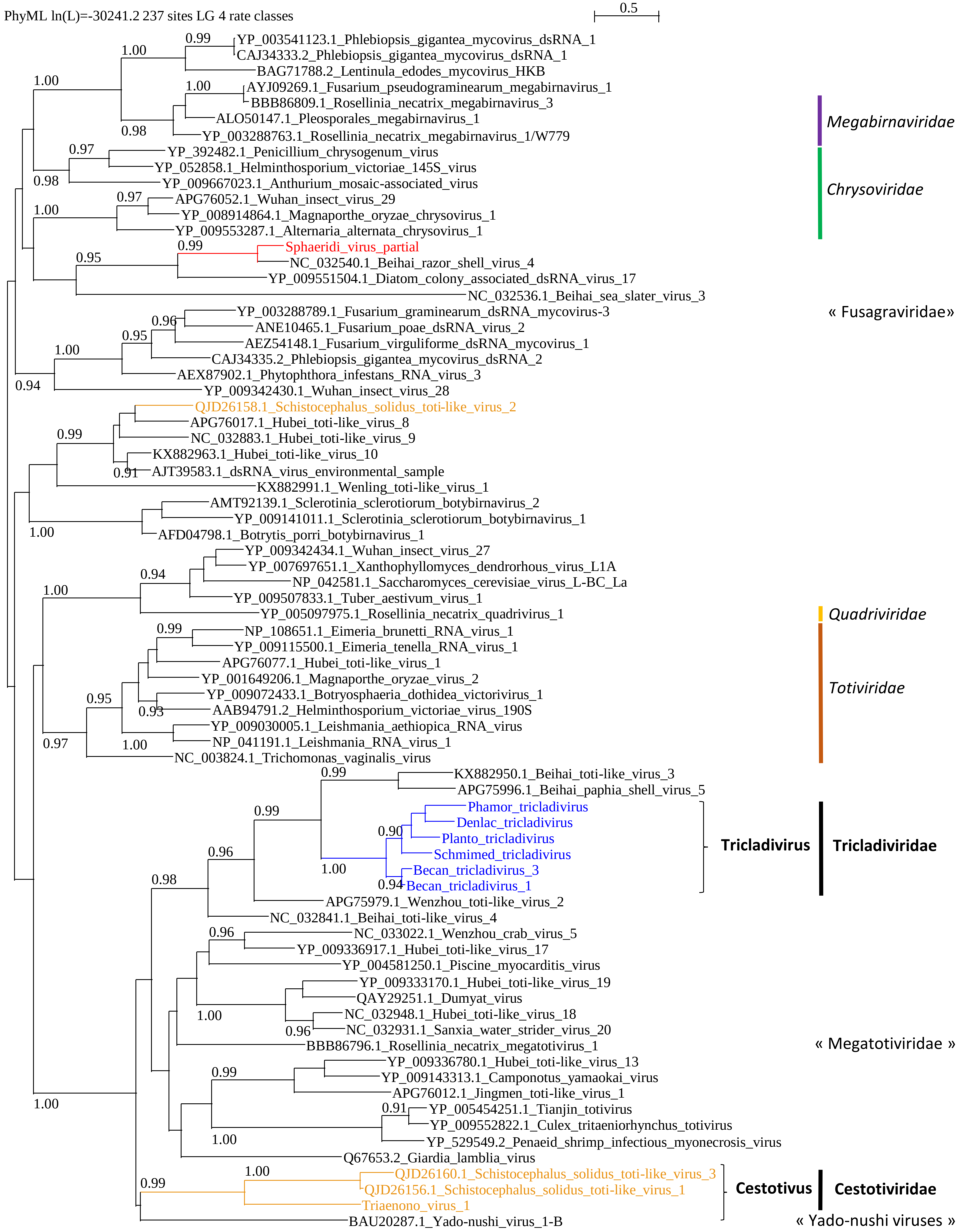


Supplementary Figure 4: Phylogenetic trees of the RNA-directed RNA polymerase (RdRPs) of RNA viruses of the order *Ghabrivirales.* Viruses of Platyhelminthes included in the trees are color coded (trematode, red; cestode, orange; Rhabditophora, blue). The trees were inferred in PhyML using the LG substitution model. Branch points indicate that results of Shimodaira-Hasgawa branch test > 0.9. Assigned family names are provided in Italic. Novel proposed genus and family names are indicated in bold and black

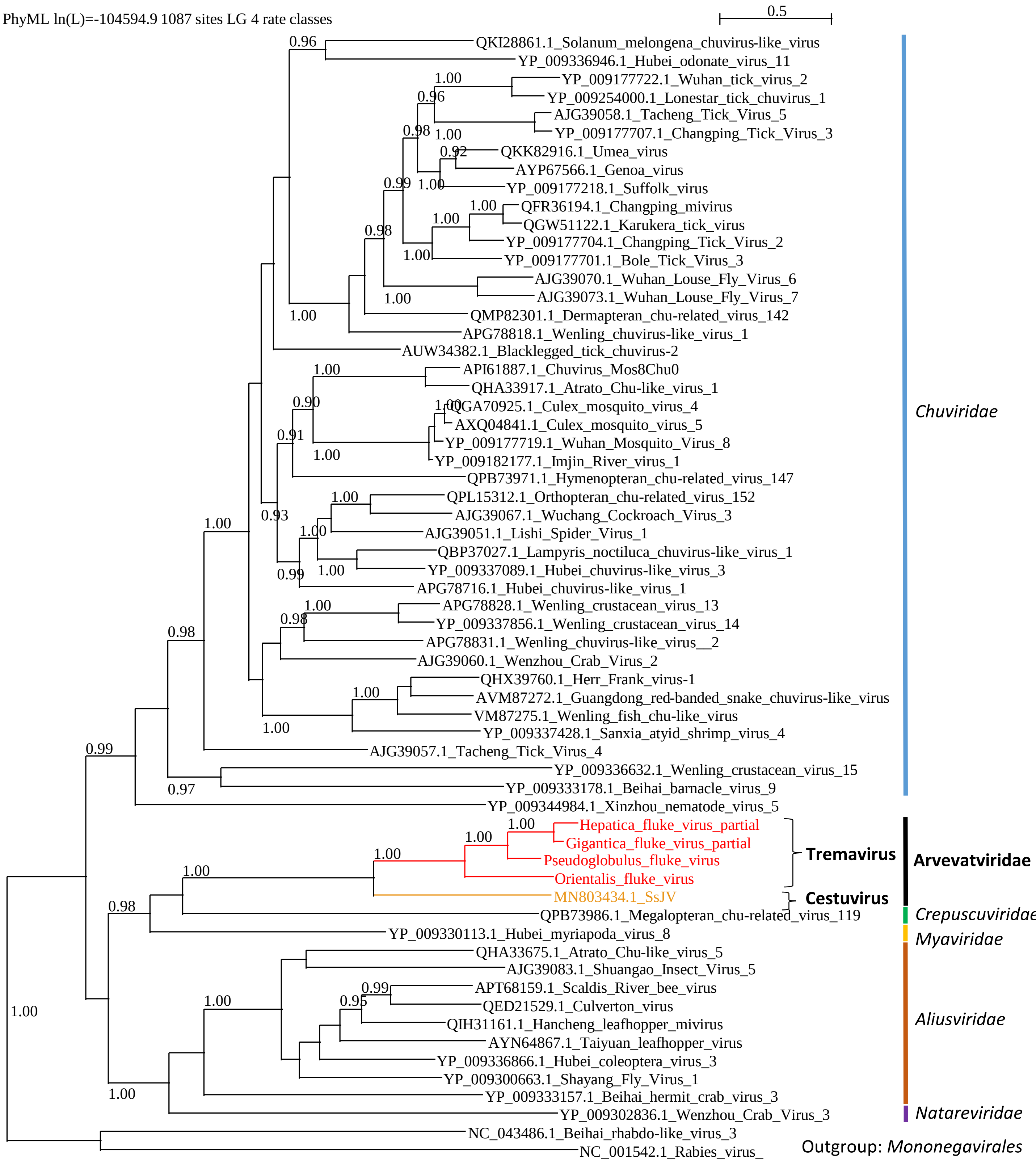


Supplementary Figure 5: Phylogenetic trees of the RNA-directed RNA polymerase (RdRPs) of RNA viruses of the order *Jingchuvirales.* Viruses of Platyhelminthes included in the trees are color coded (trematode, red; cestode, orange). The trees were inferred in PhyML using the LG substitution model. Branch points indicate that results of Shimodaira-Hasgawa branch test > 0.9. Assigned family names are provided in Italic. Novel proposed genus and family names are indicated in bold and black

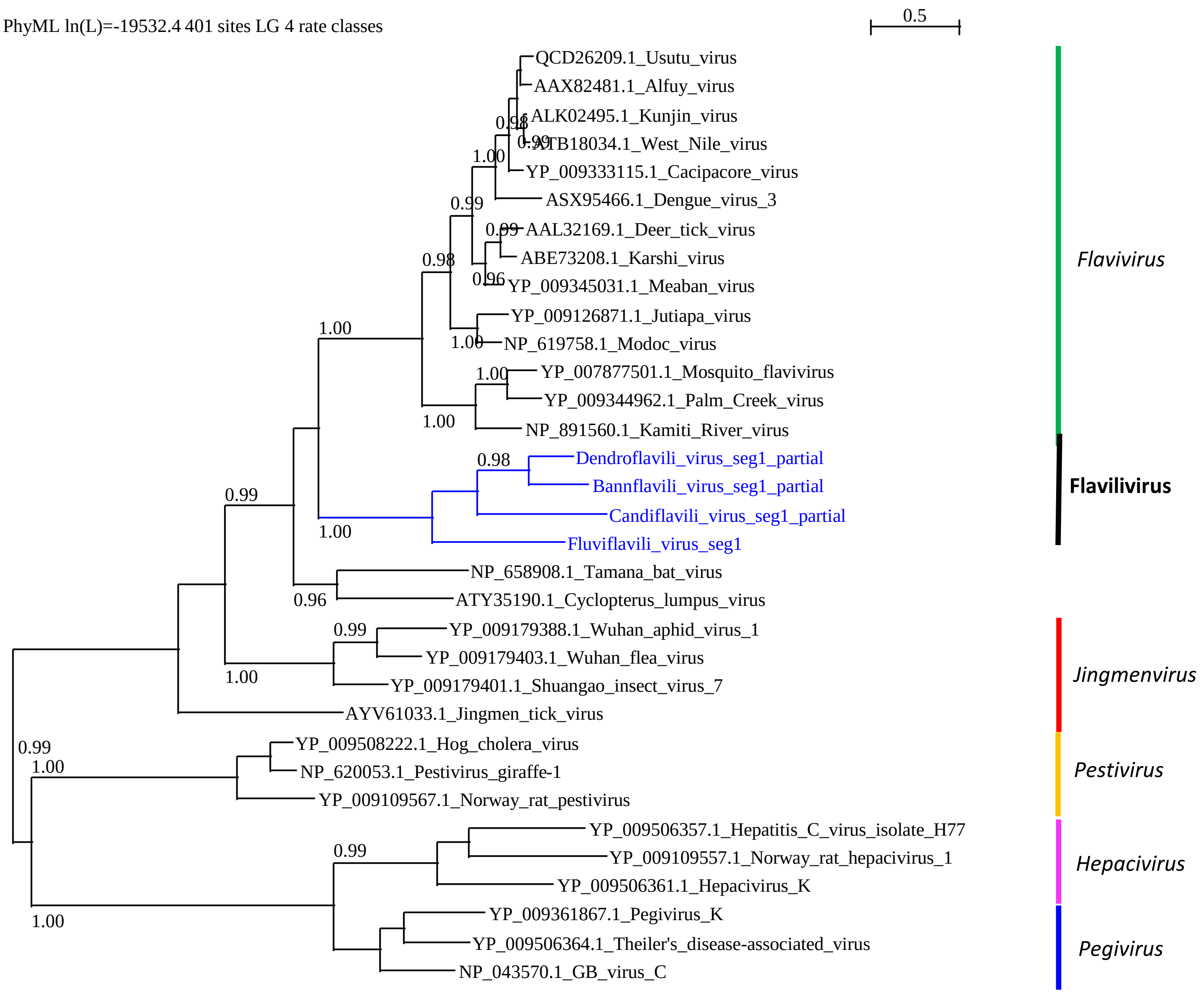


Supplementary Figure 6: Phylogenetic trees of the RNA-directed RNA polymerase (RdRPs) of RNA viruses of the family *Flaviviridae.* Viruses of Platyhelminthes included in the trees are color coded (Rhabditophora, blue). The trees were inferred in PhyML using the LG substitution model. Branch points indicate that results of Shimodaira-Hasgawa branch test > 0.9.

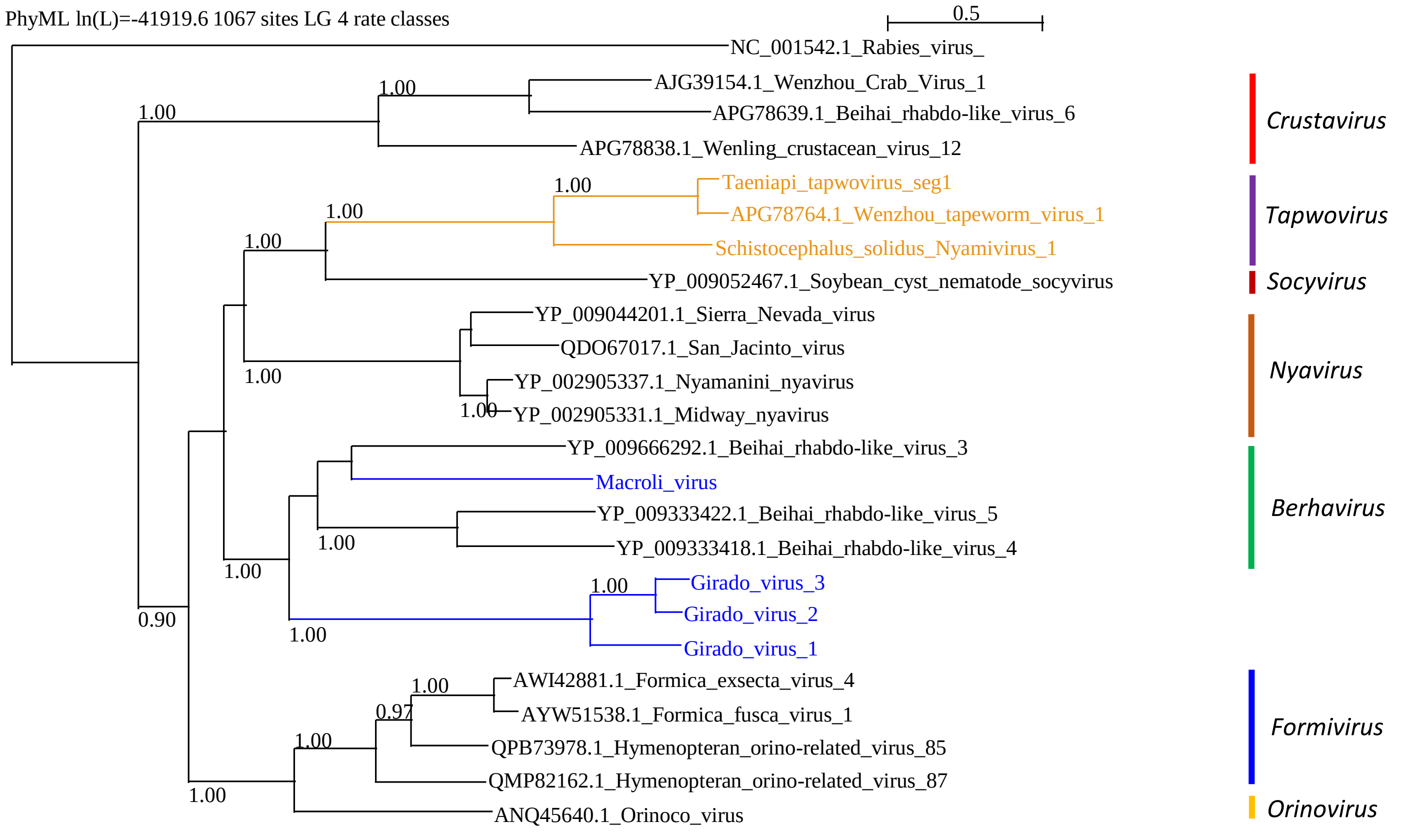


Supplementary Figure 7: Phylogenetic trees of the RNA-directed RNA polymerase (RdRPs) of RNA viruses of the family *Nyamiviridae.* Viruses of Platyhelminthes included in the trees are color coded (cestode, orange; Rhabditophora, blue). The trees were inferred in PhyML using the LG substitution model. Branch points indicate that results of Shimodaira-Hasgawa branch test > 0.9. Assigned genus names are provided in Italic.

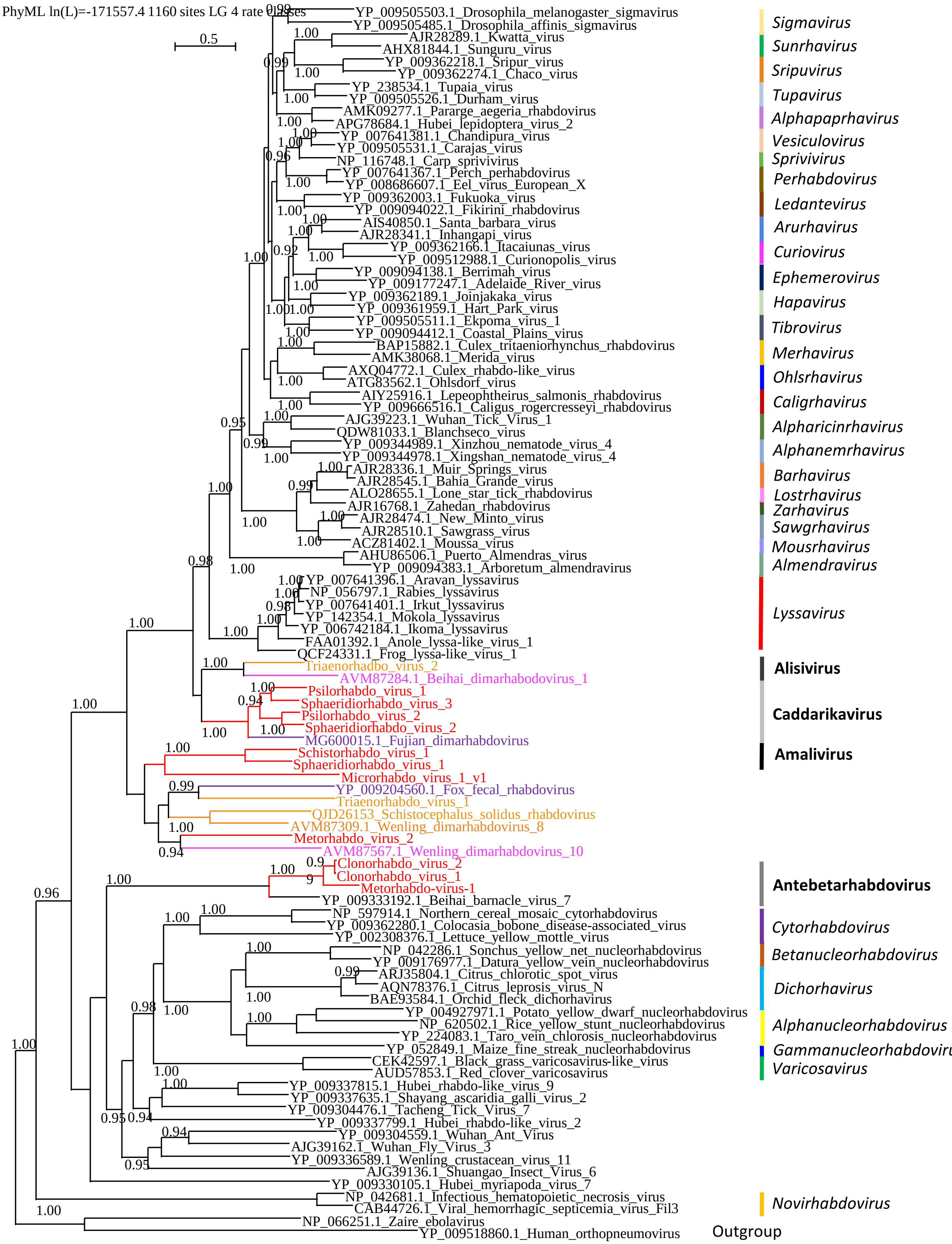


Supplementary Figure 8: Phylogenetic trees of the RNA-directed RNA polymerase (RdRPs) of RNA viruses of the family *Rhabdoviridae.* Viruses of Platyhelminthes included in the trees are color coded (trematode, red; cestode, orange; monogenean, pink; unknown Nedodermata, violet). The trees were inferred in PhyML using the LG substitution model. Branch points indicate that results of Shimodaira-Hasgawa branch test > 0.9.

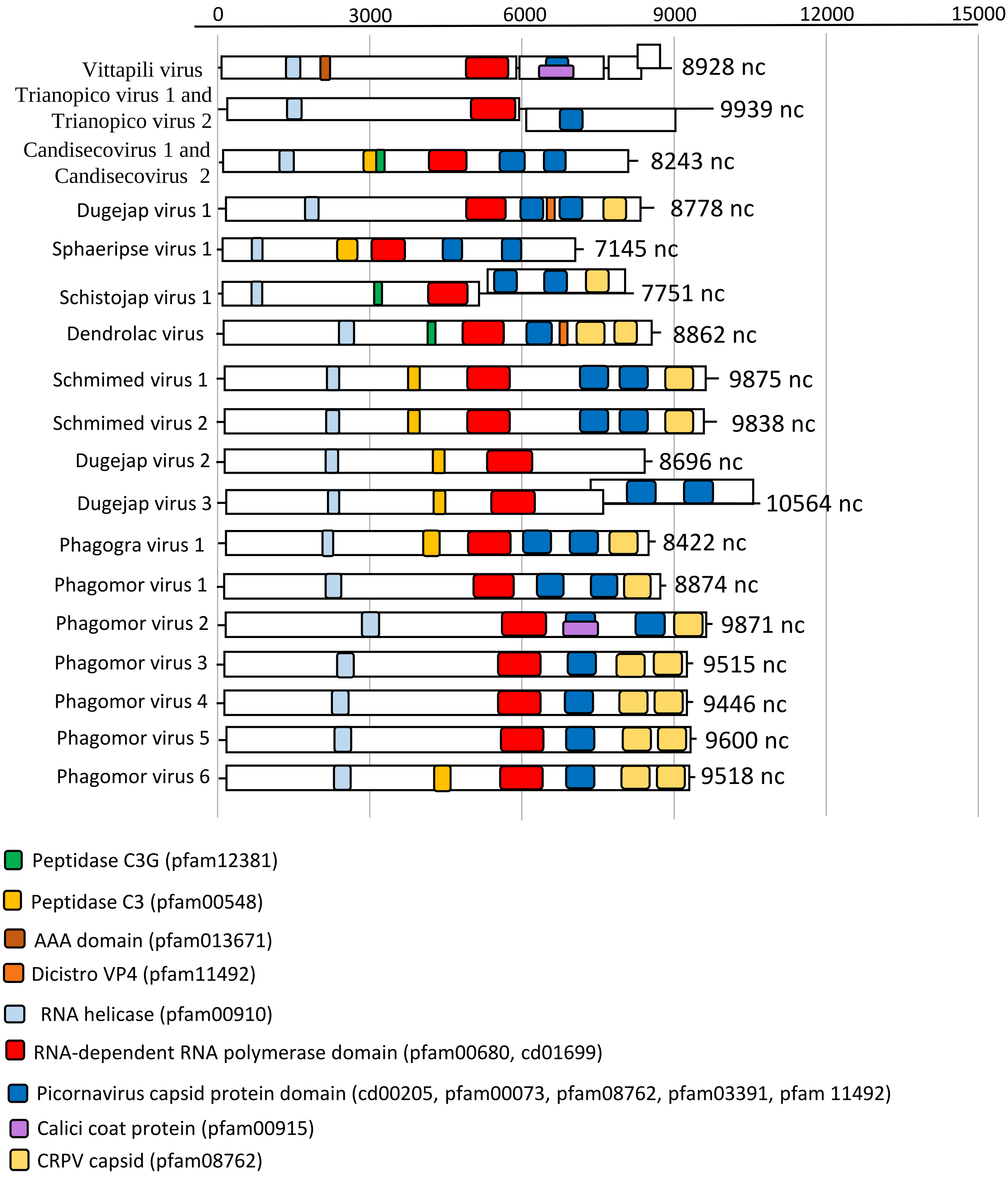


Supplementary Figure 9: Genome organization of complete sequences of RNA viruses of Platyhelminthes that belong to the order *Picornavirales.* The phylogenetic position of these viruses related to the known diversity is provided in supplementary figure 1.

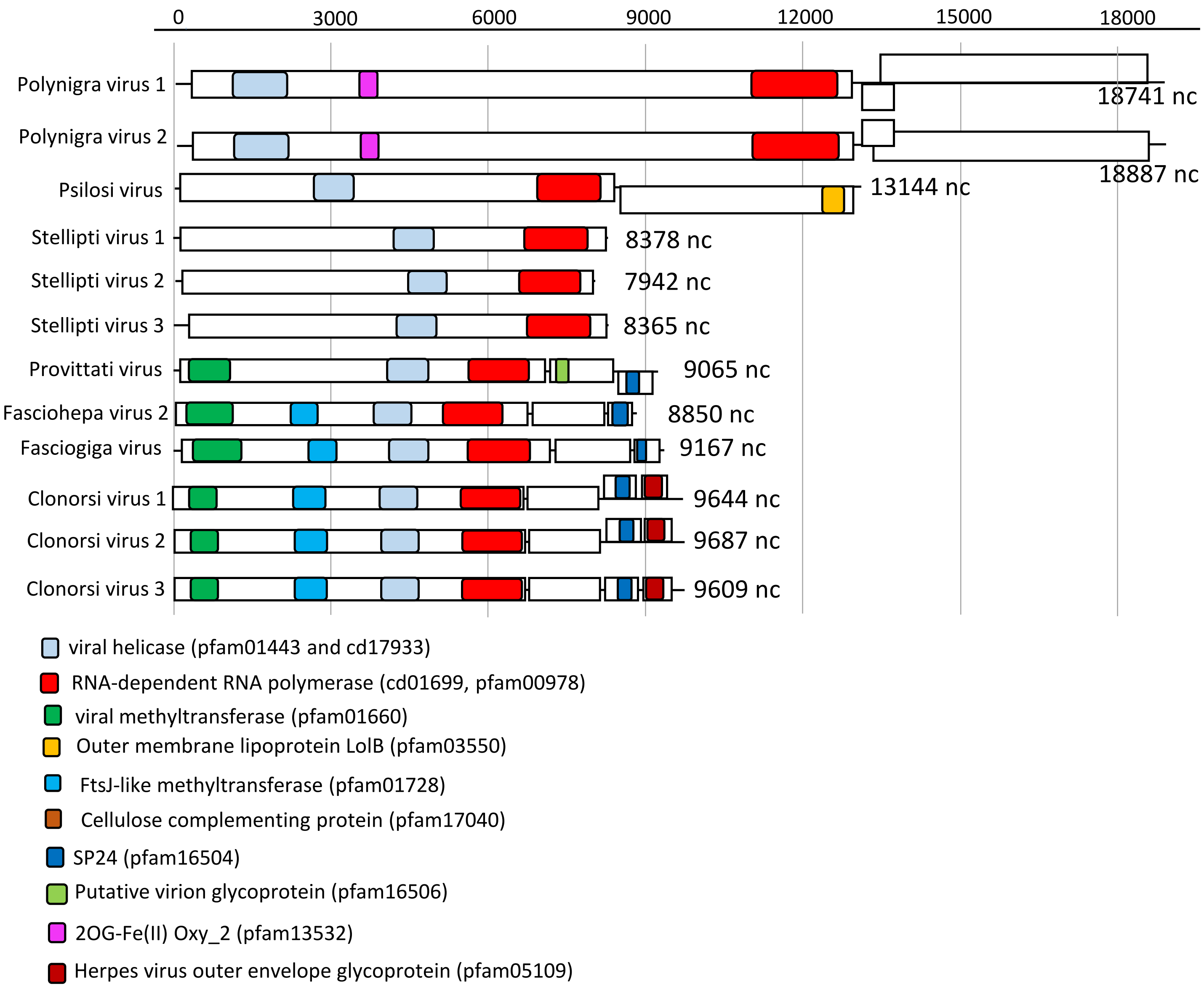


Supplementary Figure 10: Genome organization of complete sequences of RNA viruses of Platyhelminthes that belong to the order *Martellivirales.* The phylogenetic position of these viruses related to the known diversity is provided in supplementary figure 2.

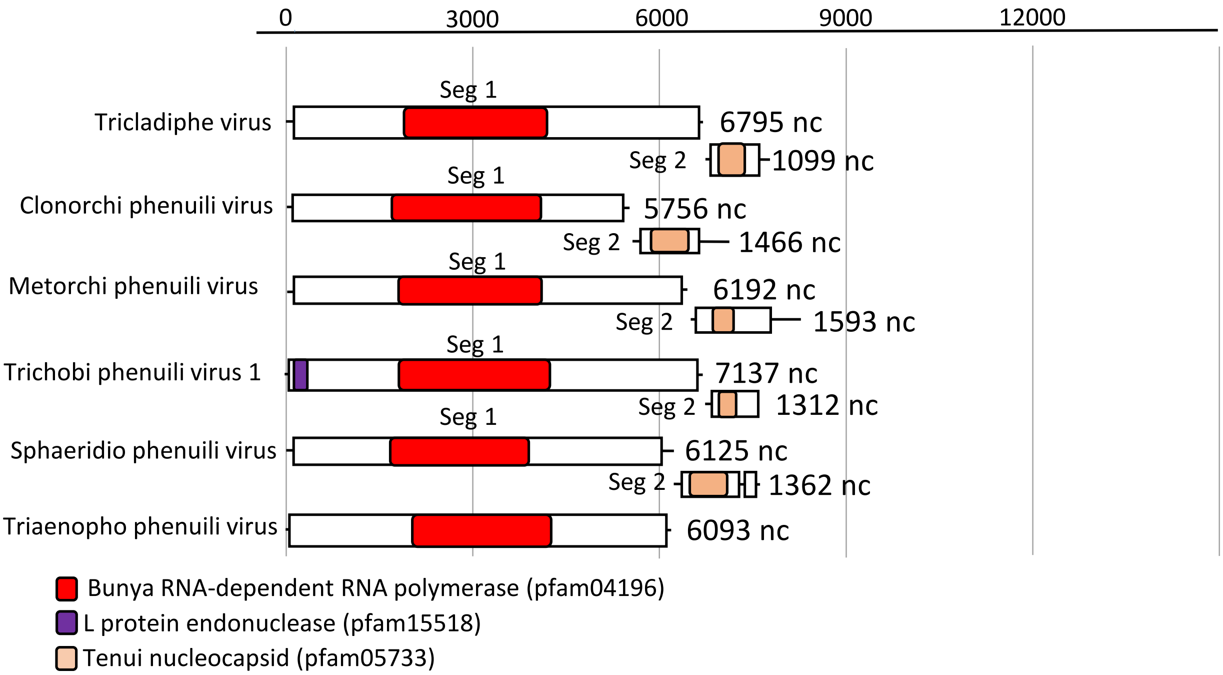


Supplementary Figure 11: Genome organization of complete sequences of RNA viruses of Platyhelminthes that belong to the order *Bunyavirales.* The phylogenetic position of these viruses related to the known diversity is provided in supplementary figure 3.

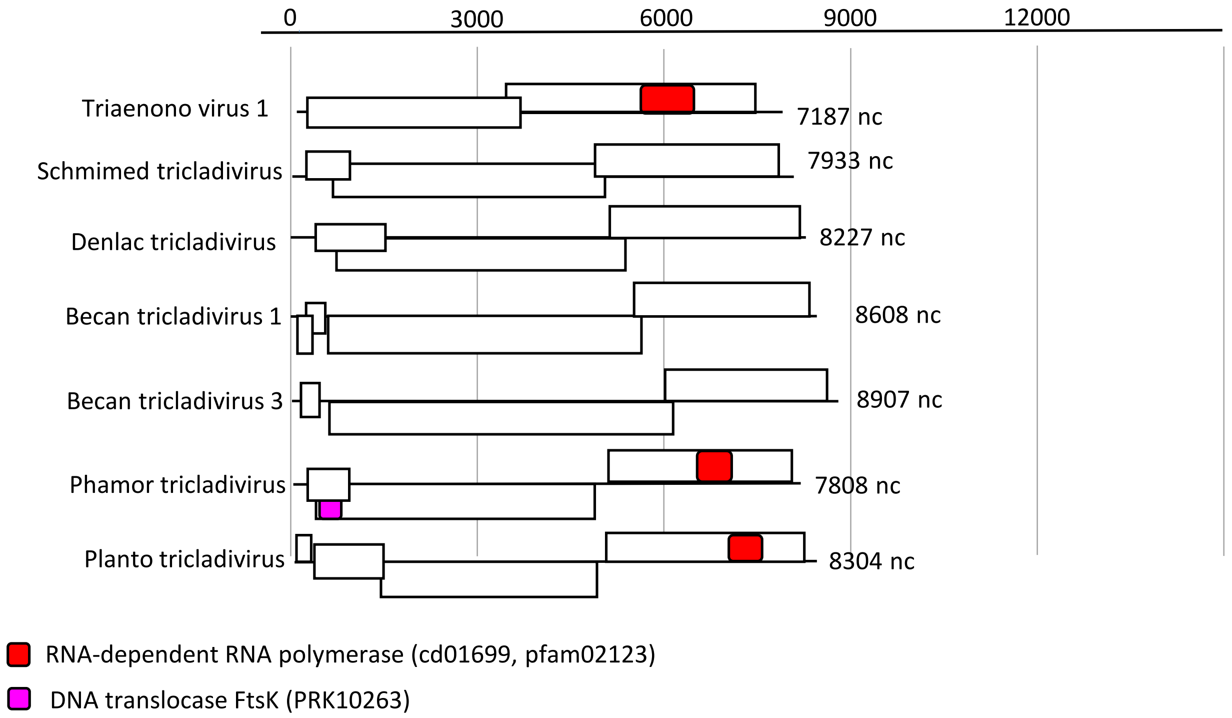


Supplementary Figure 12: Genome organization of complete sequences of RNA viruses of Platyhelminthes that belong to the order *Bunyavirales.* The phylogenetic position of these viruses related to the known diversity is provided in supplementary figure 4.

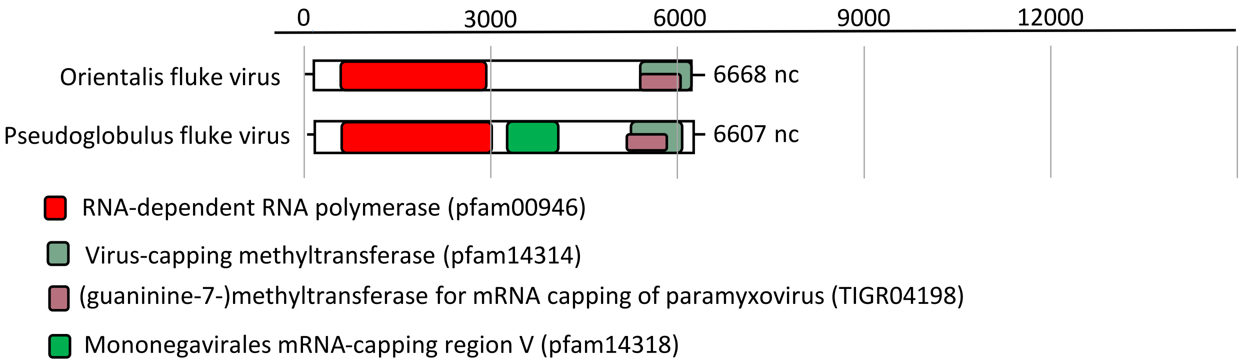


Supplementary Figure 13: Genome organization of complete sequences of RNA viruses of Platyhelminthes that belong to the order *Jingchuvirales.* The phylogenetic position of these viruses related to the known diversity is provided in supplementary figure 5.

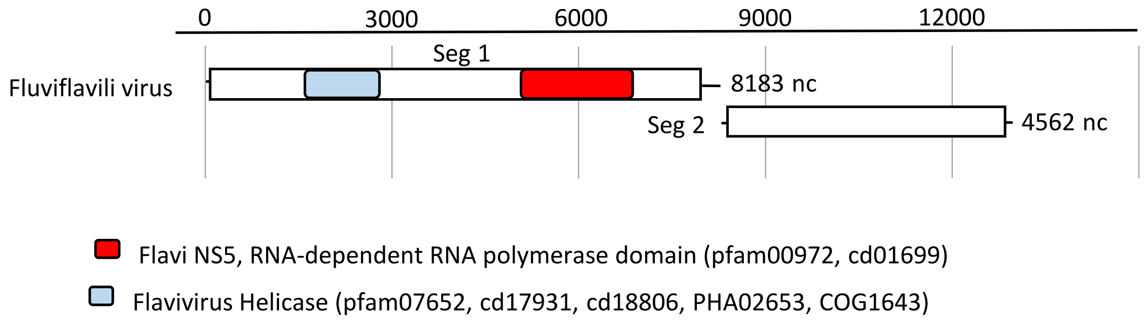


Supplementary Figure 14: Genome organization of a complete sequence of RNA viruses of Platyhelminthes that belong to the family *Flaviviridae.* The phylogenetic position of thsis virus related to the known diversity is provided in supplementary figure 6.

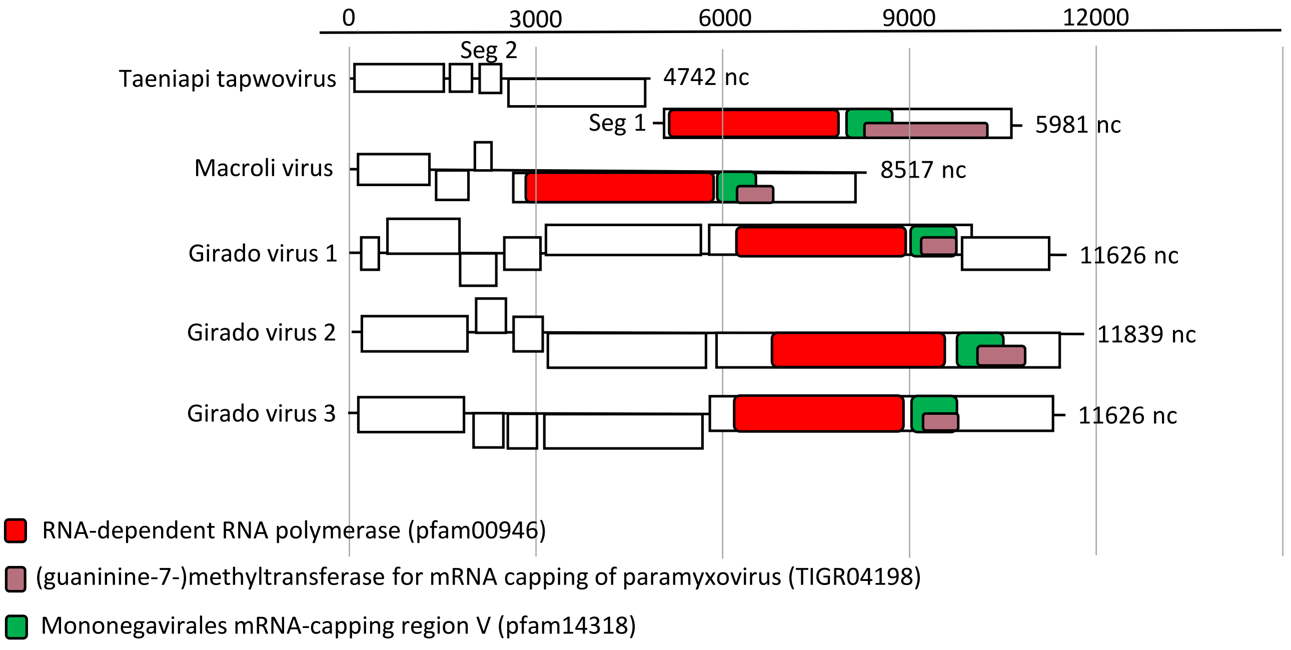


Supplementary Figure 15: Genome organization of complete sequences of RNA viruses of Platyhelminthes that belong to the family *Nyamiviridae.* The phylogenetic position of these viruses related to the known diversity is provided in supplementary figure 7.

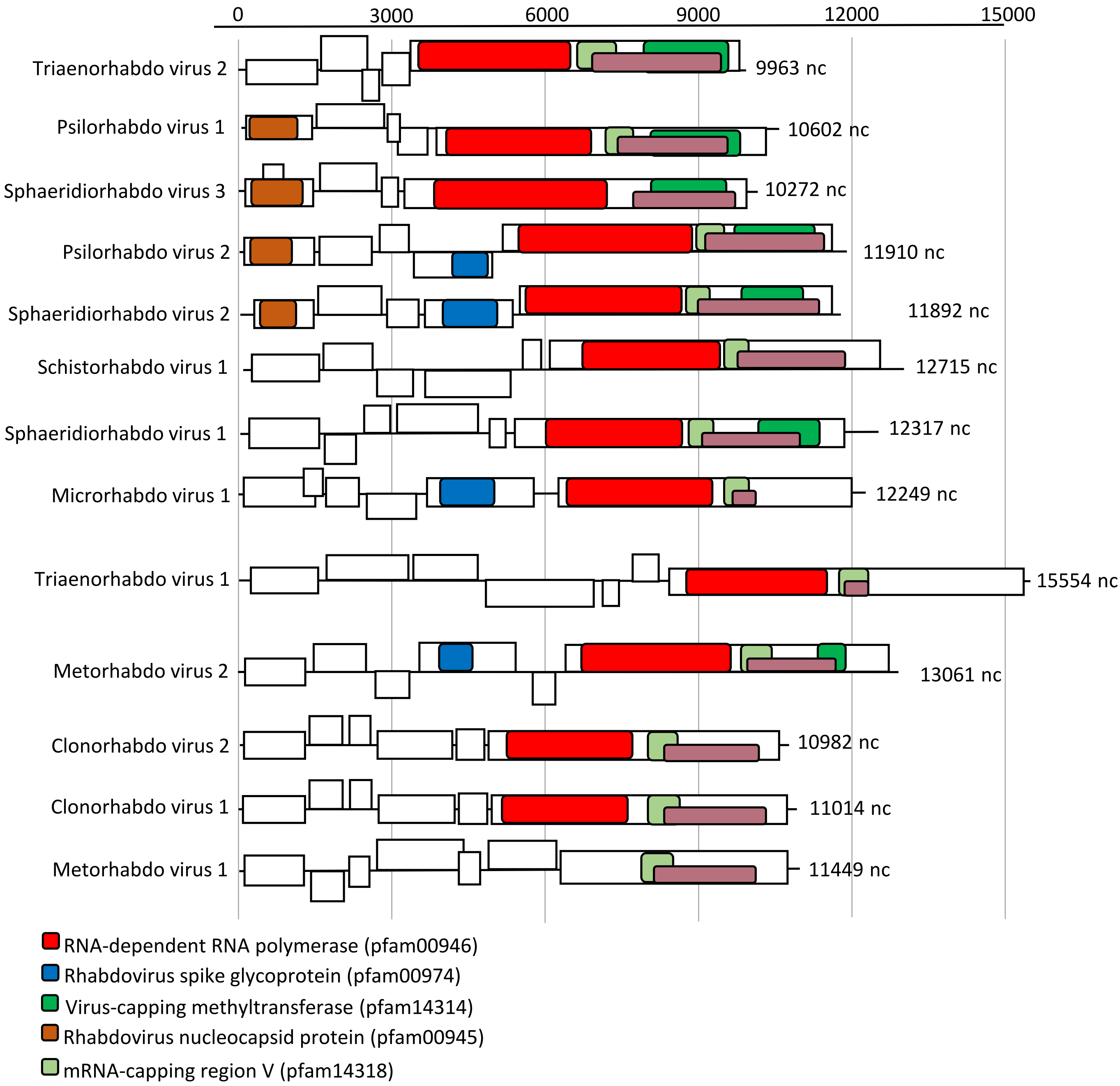


Supplementary Figure 16: Genome organization of complete sequences of RNA viruses of Platyhelminthes that belong to the family *Rhabdoviridae.* The phylogenetic position of these viruses related to the known diversity is provided in supplementary figure 8.

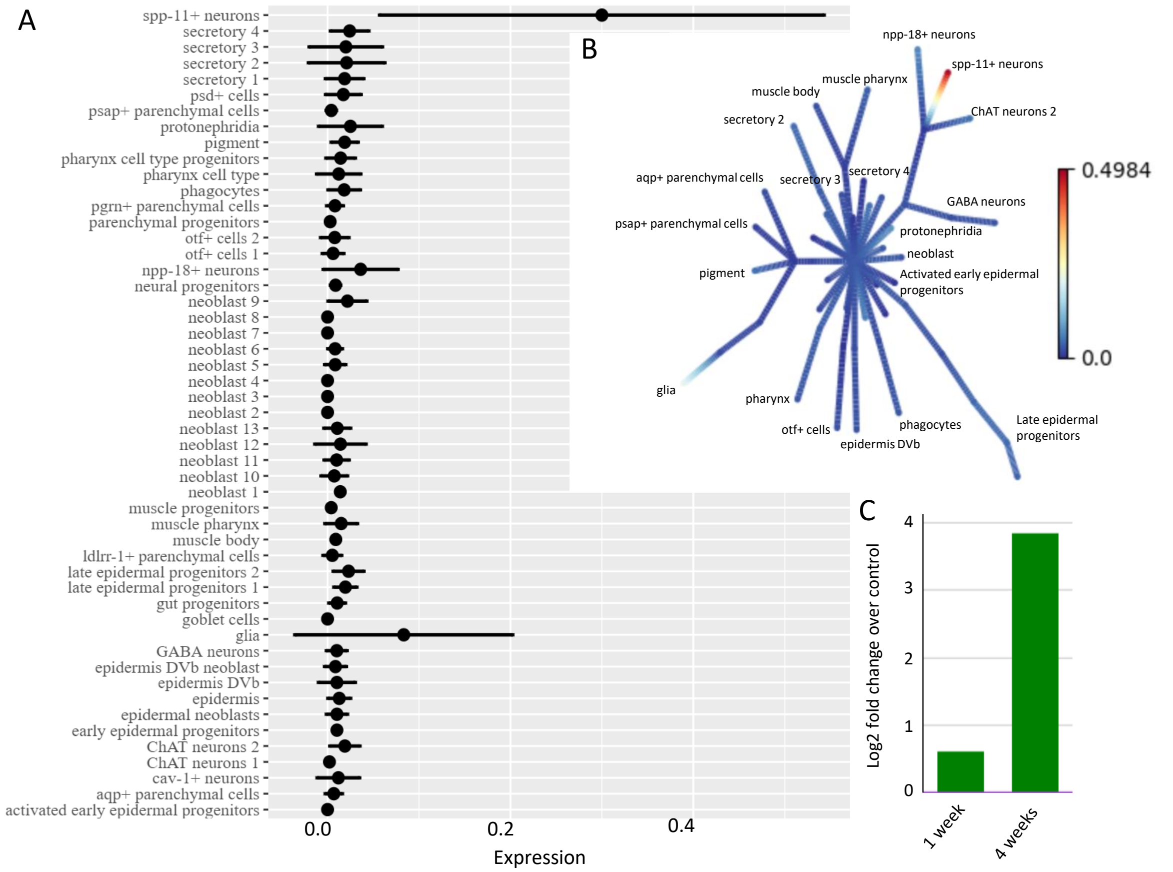


Supplementary figure 17: Schmimed virus 1 presence within *S. mediterranea* individuals used for the Cell Atlas and RNA interference studies provides clues towards cell localization and activity. A/ Schmimed virus 1 was found in spp-11+ neurons and to a lesser extent in glia, but appeared absent from other cell types identified in *S. mediterranea* single cell transcriptome atlas[54]. B/ Visualization of viral genome abundance in different cell type clusters as represented by the t-SNE pseudotime according to [54]. C/ A significant increase in Schmimed virus 1 was observed 1 week and 4 weeks following Hippo gene knockdown (as defined with edgeR with raw counts and false discovery rate cutoff of 1%) [55].
